# Components from the human c-myb transcriptional regulation system reactivate epigenetically repressed transgenes

**DOI:** 10.1101/487736

**Authors:** Cassandra M. Barrett, Reilly McCracken, Jacob Elmer, Karmella A. Haynes

## Abstract

Epigenetic silencing of transgenes has been a persistent challenge for mammalian cell engineering. Foreign DNA can be incorporated into closed chromatin before and after it has been integrated into a host cell’s genome. To identify elements that mitigate epigenetic silencing, we tested components from the c-myb and NF-kB transcriptional regulation systems in transiently transfected DNA and at chromosomally integrated transgenes in PC-3 and HEK293 cells. DNA binding sites for MYB (c-myb) placed upstream of a minimal promoter strongly enhanced expression from transiently transfected plasmid DNA. We targeted p65 and MYB fusion proteins to chromosomal transgenes that were silenced by ectopic Polycomb chromatin or by uncharacterized endogenous chromatin. Transient expression of Gal4-MYB induced sustained activation of the Polycomb-silenced *UAS-Tk-luciferase* transgene. We used custom guide RNAs and dCas9-MYB to target MYB to different sites. Transgene activation within ectopic Polycomb chromatin required proximity of dCas9-MYB to the transcriptional start site, while activation at the naturally repressed transgene was position-independent. Our report is the first to demonstrate the use of MYB in the context of the CRISPR-activation system. The results demonstrate that DNA elements and fusion proteins derived from c-myb can mitigate epigenetic silencing to improve transgene expression in engineered cell lines.

## INTRODUCTION

The advancement of cell engineering requires robust and reliable control of endogenous and synthetic genetic material within living cells. A lack of tools for enhancing the expression of transgenes in mammalian cells currently limits effective gene regulation in different contexts. Unpredictable formation of heterochromatin around transgenic material in mammalian cells limits our ability to express foreign DNA for the production of therapeutic proteins and the development of engineered mammalian systems for biosensing and computing (1, 2). Integrated transgenes are often silenced by the same mechanisms that serve as a cellular defense against viral insertion into the genome (3–5). Nucleation of heterochromatin around transgenic material can be initiated and sustained by both promoter methylation (1, 5) and various histone modifications (2, 4). For example, MyD88 pathway-mediated silencing of transgenes leads to an accumulation of repressive H3K9me on newly bound histones (2, 6). Silencing of transgenes may also be Polycomb-mediated, where Polycomb repressive complexes deposit H3K27me3 on histones to establish a silenced state (7–9). The diversity and persistence of transgene silencing has led to the development of tools for mammalian cell engineering specifically aimed at combating heterochromatin.

Recruiting activators to a specific locus in order to reverse epigenetic silencing can be achieved either by including an activation-associated cis-regulatory DNA sequence within the construct itself, or through the targeting of engineered fusion proteins to the silenced transgene. Both natural and synthetic cis-regulatory motifs that recruit activators have been used (10–13) to help increase transgene expression as an alternative to viral promoters that are prone to methylation and silencing (1). Previous screens by ourselves and other groups (11, 14, 15) have identified mammalian activation-associated cis-regulatory elements that recruit endogenous factors to increase the expression of epigenetically silenced transgenes, including motifs for nuclear factor Y, CTCF, and elongation factor alpha (EF1-α) (12, 13). The underlying regulatory mechanisms are not entirely understood, since in this case efficient screening for functional sequences has been prioritized over dissecting the mechanism of individual elements.

Fusion proteins that target activation-associated domains to transgenes can also be used to reverse silencing. Targeted epigenetic effectors such as p300 (histone acetyltransferase) and Tet1 (methylcytosine dioxygenase) are potent activators of gene expression. (16–18) These directly alter local chromatin features, therefore their function may be context dependent. (17, 18) Transcriptional activation domain (TAD) peptides, including Herpes simplex virus protein vmw65 tetramer (4x VP16, VP64) and nuclear factor NF-kappa-B p65 subunit (p65) have been used singly, or as subunits within compound activators such as VPR, SAM, and SunTag (19–21). Site-specific targeting of VP64 (4x VP16) enhances endogenous gene expression, and remodels chromatin through the accumulation of activation-associated histone modifications (H3K27ac and H3K4me) (20, 22, 23). p65-based systems are also very effective at restoring both endogenous (19, 24) and transgenic (25) gene expression.

Significant progress towards transgene reactivation has been made so far, but several important gaps remain. First, several natural mechanisms of activation are not yet represented in published cell engineering studies. Chromatin remodelers that shift, remove, or exchange nucleosomes (26), and pioneer factors increase DNA accessibility in closed chromatin by displacing linker histones (26–28) remain under-utilized for transgene regulation. Second, the critical parameters for stable transgene activation are not yet fully defined. So far, at least two studies have demonstrated prolonged enhancement of transgenes (10 to 25 days) via targeted fusion proteins alone (29) or in combination with flanking anti-repressor DNA elements (30). Neither study evaluated the chromatin features at the target genes prior to their reactivation, therefore the context in which expression enhancement occurred is uncertain. Finally, the performance of targeted activators can be context-dependent. Catalytic domains used for site-specific chromatin remodeling (18, 30, 31), may be inhibited by pre-existing chromatin features that vary across loci. For example, Cano-Rodriguez *et al.* constructed a targeted histone methyltransferase fusion and found that the endogenous chromatin microenvironment, including DNA methylation and H3K79me, impacted the ability of their fusion protein to deposit H3K4me and induce activation (32). Similarly inconsistent performance has been shown for other fusions that generate H3K79me and H3K9me (33, 34). Systematic studies at loci with well-defined chromatin compositions are needed to fully understand mechanisms of chromatin state switching.

Here, we expand our previous work where we had identified cis-regulatory sequences that enhanced expression from plasmid-borne transgenes (12). To regulate expression of chromosomally-inserted transgenes, we built site-specific fusion proteins with effector modules that represent diverse activities: transcriptional activation through cofactor recruitment, direct histone modification, and nucleosome repositioning and displacement. We focus on reversal of silencing within Polycomb heterochromatin, which is known to accumulate at transgenes that are integrated into chromosomes (7–9) and is widely distributed across hundreds or thousands of endogenous mammalian genes that play critical roles in normal development and disease (9, 35, 36). We report that recruitment of p65 and MYB-associated components via a cis-regulatory element or fusion proteins enhances expression from epigenetically silenced transgenes. MYB-mediated activation within Polycomb heterochromatin relies on interactions with p300 and CBP. Our results have implications for determining the most appropriate strategy to enhance gene expression, specifically within Polycomb-repressed chromatin.

## MATERIALS AND METHODS

### Construction and Testing of Plasmids containing MYB- and p65 Motifs

Plasmid construction, transfection of PC-3 cells, and luciferase assays were carried out as described previously (12). Briefly, cloning of double-stranded oligos was used to insert motifs 222 bp upstream of the transcription start site of an EF1a promoter at XbaI/SphI. Plasmids were then transfected into PC-3 cells (ATCC, CRL-1435) using Lipofectamine LTX™ following the manufacturer’s recommended protocols. Luciferase expression was measured 48 hours after transfection using a luciferase assay kit (Promega, Madison, WI). All luciferase values were normalized relative to the native plasmid control, which contained an unaltered EF1a promoter.

### Construction of MV14 and Gal4-AAP Plasmids

We constructed mammalian expression vector 14 (MV14) for the overexpression of Gal4-mCherry-AAP fusion proteins in-frame with a nuclear localization sequence and 6X-histidine tag. First, plasmid MV13 was built by inserting a Gal4-mCherry fragment into MV10 (37) directly downstream of the CMV promoter. Next, MV14 was built by inserting a SpeI/PstlI (FastDigest enzymes, ThermoFisher Scientific) -digested gBlock Gene Fragment (Integrated DNA Technologies), which encoded a XbaI/NotI multiple cloning site, into MV13 downstream of mCherry. Ligation reactions included gel-purified (Sigma NA1111) DNA (25 ng linearized vector, a 2x molar ratio of insert fragments), 1x Roche RaPID ligation buffer, 1.0 uL T4 ligase (New England Biolabs), in a final volume of 10uL.

AAPs were cloned into MV14 at the multiple cloning site containing XbaI and NotI cut sites. AAPs were either ordered from DNASU in vectors and amplified using primers that added a 5’ XbaI site and a 3’ NotI site or ordered as gBlock Gene Fragments with the same 5’ and 3’ cutsites (Integrated DNA Technologies). Sequences in vectors were amplified with Phusion High Fidelity DNA Polymerase (New England BioLabs) and primers listed in Supplementary Table S2. MV14 and AAP inserts were double-digested with FastDigest *XbaI* and FastDigest *NotI* (ThermoFisher Scientific) and then ligated with T4 DNA ligase (New England Biolabs). MV14_AAP plasmids are publicly available through DNASU (Supplementary Table S4)

### Cell Culturing and Transfections

Luc14 and Gal4-EED/luc HEK293 cells were grown in Gibco DMEM high glucose 1× (Life Technologies) with 10% Tet-free Fetal Bovine Serum (FBS) (Omega Scientific), 1% penicillin streptomycin (ATCC) at 37 °C in a humidified 5% CO_2_ incubator. Gal4-EED/luc cells were treated with 1 µg/mL doxycycline (Santa Cruz Biotechnology) for 2 days to induce stable polycomb repression. Dox was removed and cells were cultured for another four days before being seeded in 12-well plates. Luc14 cells and dox-induced Gal4-EED/luc cells were seeded in 12-well plates such that cells reached 90% confluency for lipid-mediated transfection. Transfections were performed with 1 µg plasmid per well, 3 µL Lipofectamine LTX, and 1 µL Plus Reagent (Life Technologies) per the manufacturer’s protocol. Seventy-two hours post transfection, cells were either collected for analysis or passaged further.

Puromycin selection was carried out on Gal4-AAP-expressing cells for the experiments represented in Figure 5 and Supplemental Figure S1. Dox-treated Gal4-EED/luc cells were transfected in 12-well plates and then grown for 24 hours before the addition of 10 µg/mL puromycin (Santa Cruz Biotechnology) to Gibco DMEM high glucose 1× (Life Technologies) with 10% Tet-free Fetal Bovine Serum (FBS) (Omega Scientific), 1% penicillin streptomycin (ATCC). Cells were grown in puromycin containing media for two days before wash out.

**Figure 1.**
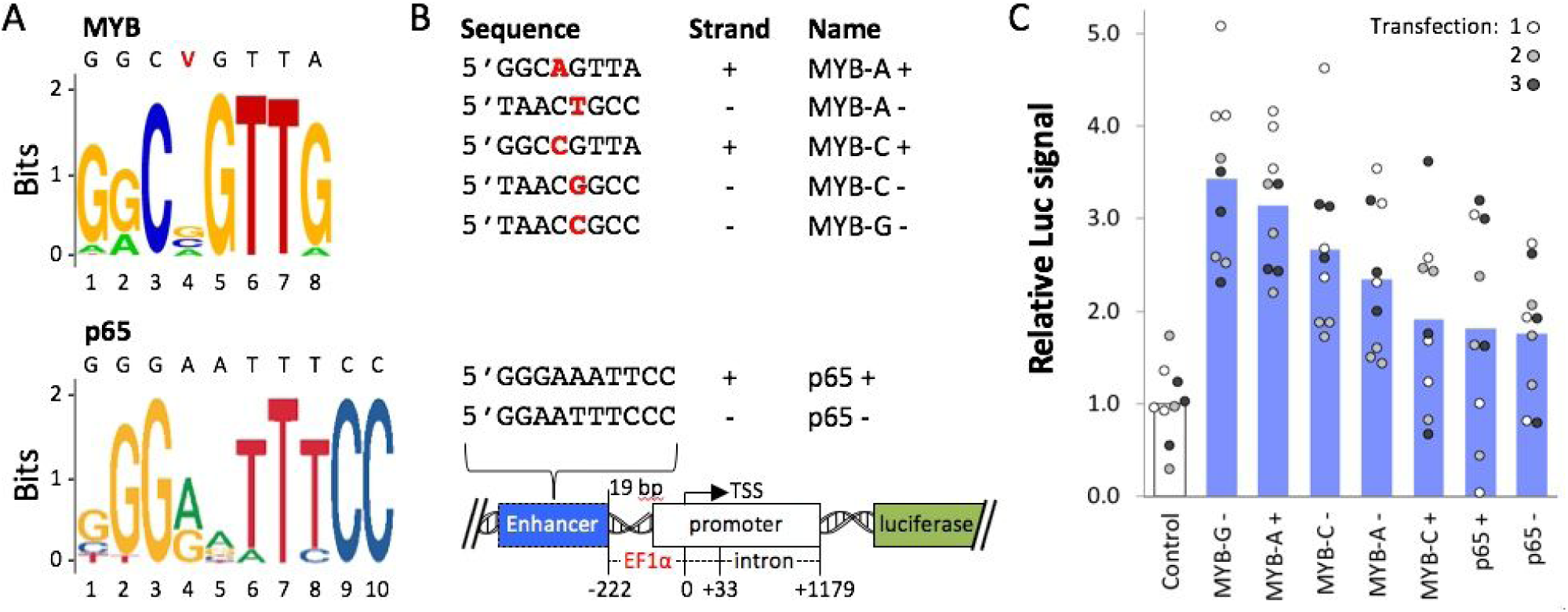
Luciferase expression from MYB- and p65-enhancer constructs. (**A**) Enhancer motif logos for MYB and p65 were generated by JASPAR (60). The MYB sequence includes a variable site (V) equally represented by A, C, or G nucleotides. (**B**) *Luciferase* reporter constructs included one of the enhancer sequences (MYB-A+, etc.) 19 bp upstream of an EF1α promoter, or no enhancer (Control). (**C**) Luciferase assays were carried out using PC-3 cells transfected with Lipofectamine-plasmid complexes. For each transfection, luminescence (Luc signal) values were measured in triplicate and normalized to the average signal from the Control. Circle = one Luc measurement, bar = mean of nine Luc values.

**Figure 2.**
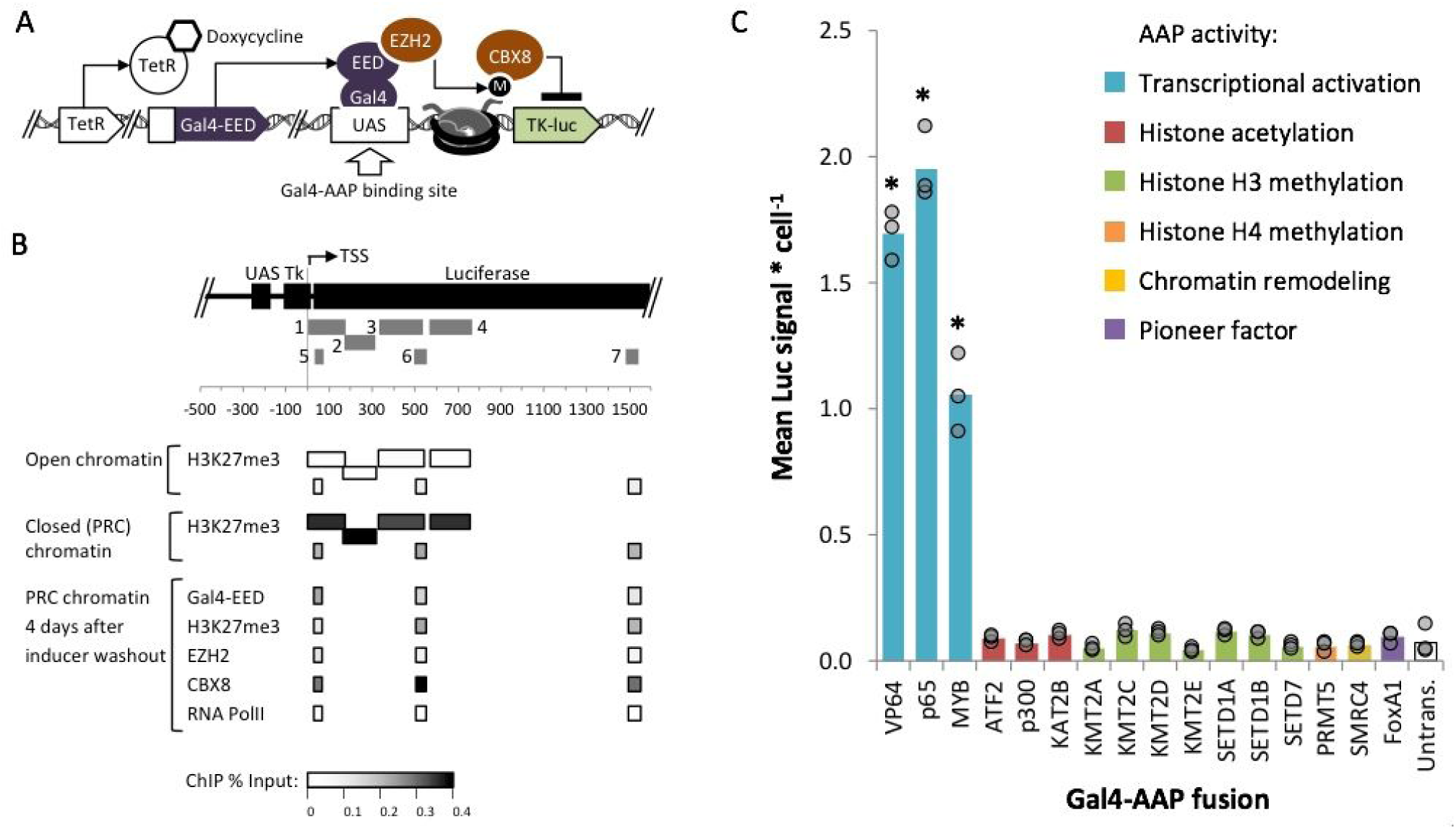
Measurement of *luciferase* reporter expression within closed or open chromatin after exposure to Gal4-AAP fusions. (**A**) In Gal4-EED/luc HEK293 cells expression of the Gal4-EED fusion protein is controlled by a Tetracycline repressor (TetR). Treatment with dox allows expression of Gal4-EED, which binds UAS and recruits EZH2 (a subunit of PRC2). EZH2 methylates (M) histone H3K27, which recruits CBX8 (a subunit of PRC1) (**B**) Panel B summarizes published chromatin immunoprecipitation (ChIP) data from previous analyses of the *Tk-luciferase* locus. Grey numbered bars indicate amplicons for quantitative PCR: 1-4 (25), and 5-7 (68). (**C**) Dox-treated cells were transfected with each Gal4-AAP fusion plasmid. Three days after transfection luciferase signal was measured. Each circle in the bar graph shows the mean luciferase (Luc) signal for a single transfection, divided by cell density (total DNA, Hoechst staining signal). Bars show means of three transfections. Asterisks (*) = *p* < 0.05 compared to untransfected cells.

**Figure 3.**
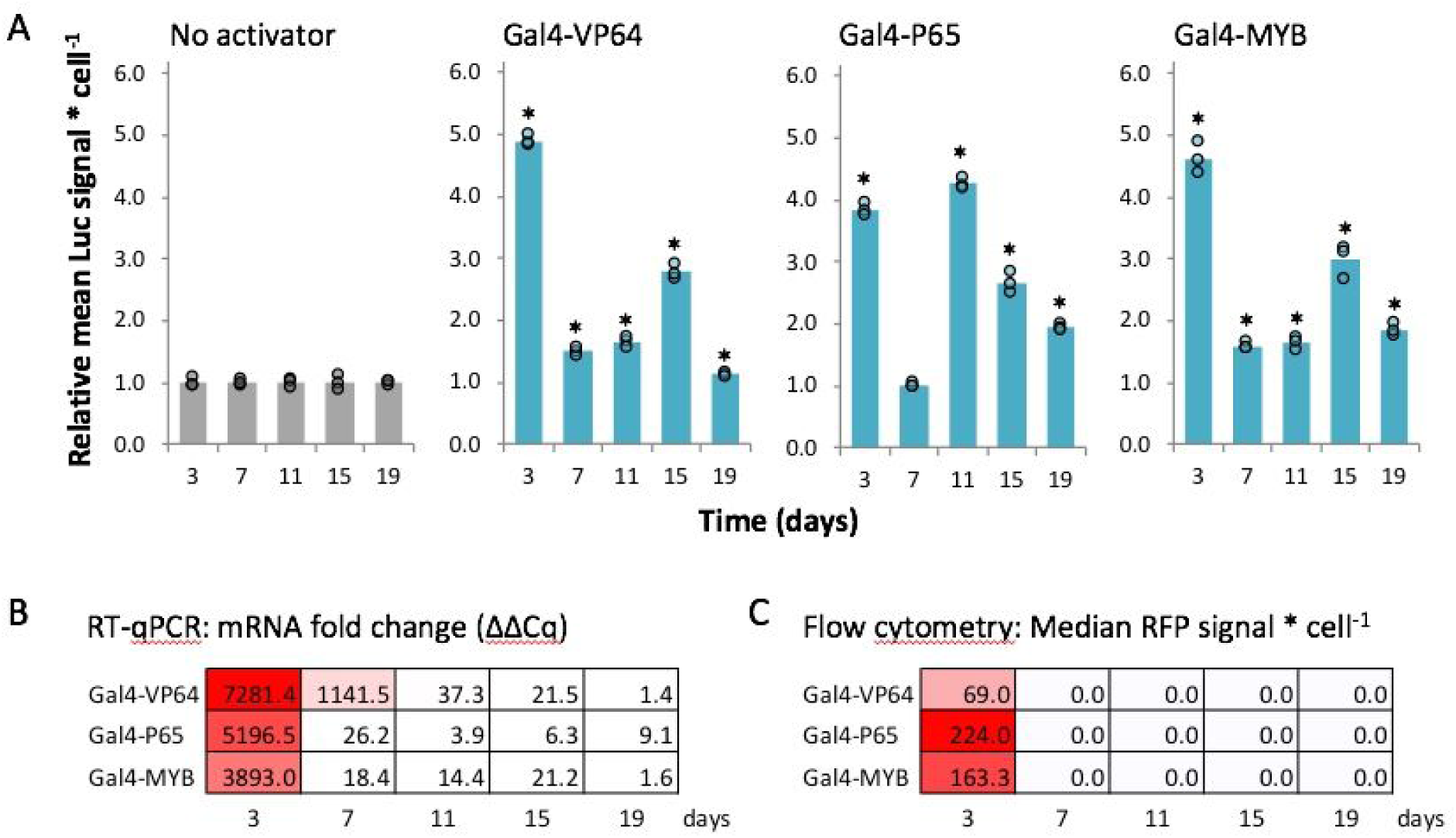
Expression of Polycomb-repressed *Tk-luciferase* over time after expression and loss of Gal4-P65, Gal4-VP64 or Gal4-MYB. (**A**) Gal4-EED/luc cells were treated with dox to induce polycomb chromatin, transfected with a Gal4-AAP plasmid, and grown under puromycin selection (10 μg/mL) for two days. At 3 days after selection, cells were sampled for luciferase (Luc) assays, passaged in puromycin-free medium, then sampled 7, 11, 15, and 19 days post selection for additional Luc assays. Mean Luc signal per cell is presented as described for Figure 4, except individual values (circles) at each time point are normalized by the mean of the “No activator” negative control. Asterisks (*) = *p* < 0.05 compared with the negative control. Results from an additional trial are shown in Supplementary Figure S4. (**B**) Reverse transcription followed by quantitative PCR (RT-qPCR) with primers for the universal mCherry region was used to determine Gal4-AAP transcript levels. “mRNA fold change” represents the Cq value normalized by the Cq of a housekeeping gene (*TBP*), and relative to untransfected “No activator” cells (Lipofectamine reagent only), log2 transformed. (**C**) Flow cytometry of mCherry signal (red fluorescent protein, RFP) was used to determine Gal4-AAP protein levels. Data in B and C were generated from one set of transfections in A. For other samples, cells were visually inspected for RFP to verify the loss of Gal4-AAP.

**Figure 4.**
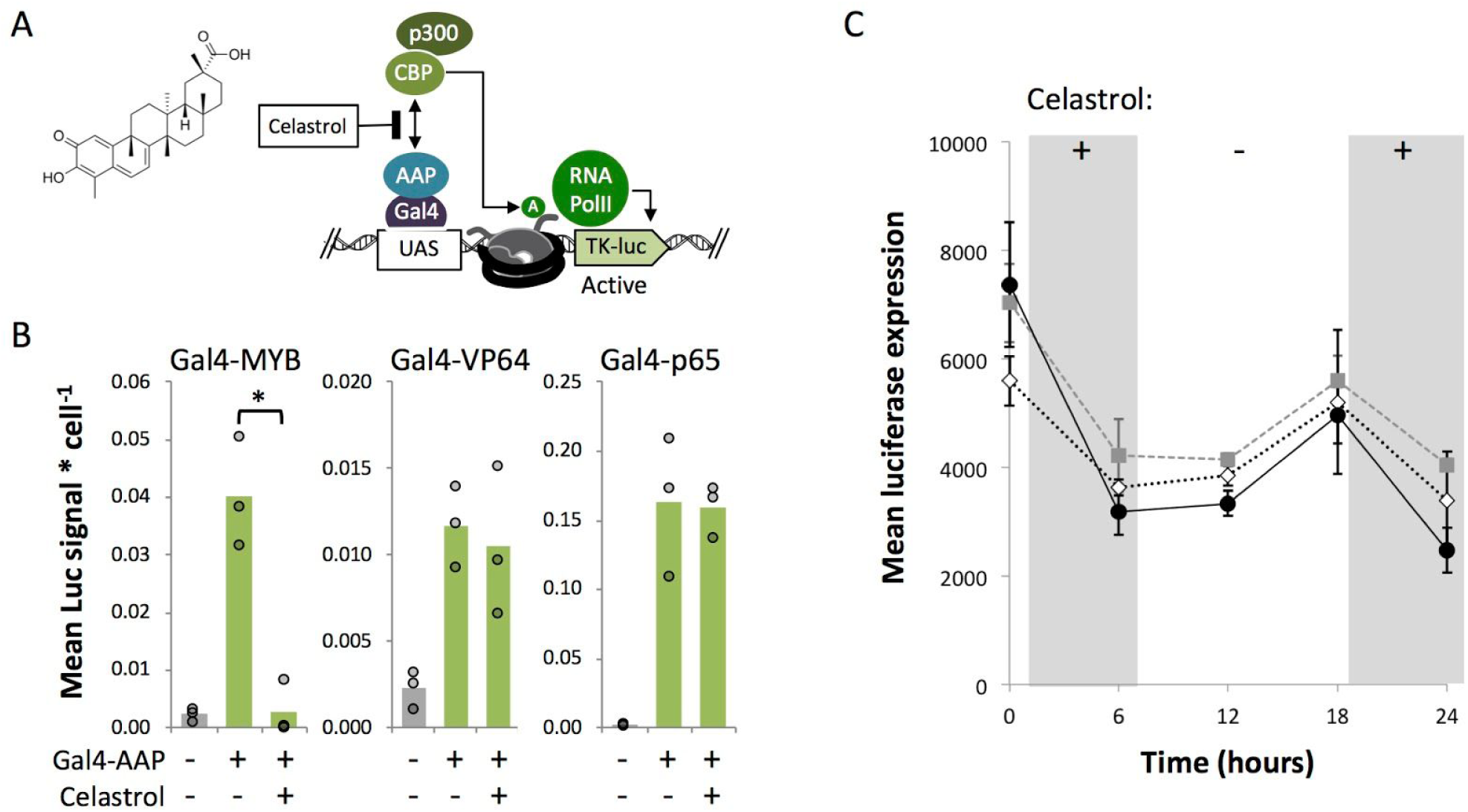
Celastrol disrupts Gal4-MYB-mediated activation of *luciferase* in closed chromatin. (**A**) The p300/CBP complex acetylates histones via the catalytic HAT domain of p300 and/or CBP (73). Celastrol inhibits the recruitment of p300/CBP by MYB by binding a docking domain in CBP that facilities complex assembly (75, 77). (**B**) Three days after Gal4-EED-mediated repression of *Tk-luciferase* and transfection with Gal4-AAPs, cells were treated with 5 μM celastrol for six hours and collected for luciferase assays. Mean luciferase (Luc) signal per cell is presented as described for Figure 3. Asterisk (*) = *p* < 0.05. (**C**) Luc measurements were carried out in Gal4-MYB-expressing cells after removal (-) and re-addition (+) of celastrol. Each series represents an independent transfection. Point = mean of three luciferase assays, bars = standard error.

**Figure 5.**
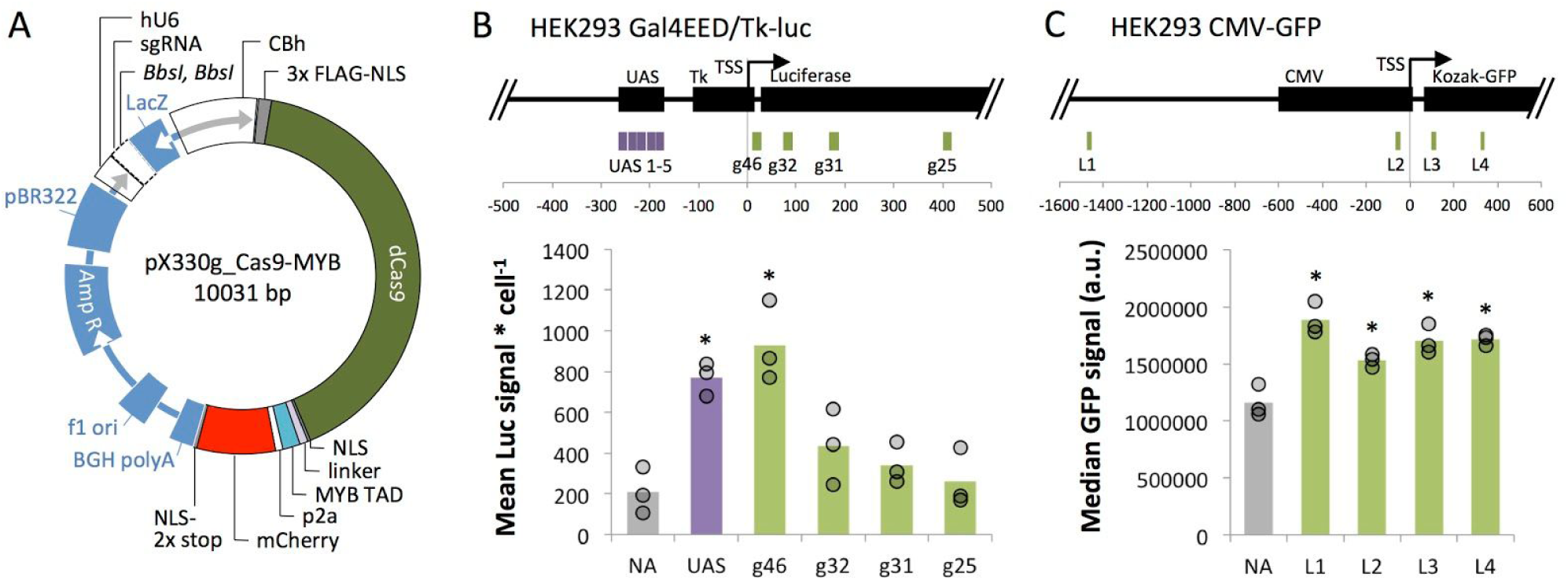
dCas9-MYB’s ability to enhance expression in induced Polycomb heterochromatin is dependent upon distance from the promoter. (**A**) Expression vector pX330g_dCas9-MYB was constructed from vector pX330A_dCas9 (a gift from Takashi Yamamoto, Addgene plasmid #63598) to co-express a dCas9-MYB fusion protein and mCherry from a CBh promoter. Single-stranded guide RNA sequences (Supplementary Table S4) were cloned into the BbsI sites and expressed from a hU6 promoter on the same vector. (**B**) We targeted dCas9-MYB to four locations (g46, g32, g31, g25) across the *Tk-luciferase* transgene in silenced Gal4-EED/luc cells. Mean luciferase signal per cell is presented as described for Figure 4. The control (grey bar) represented untransfected cells treated with Lipofectamine only (No Activator, NA). (**C**) We targeted dCas9-MYB to four sites (L1-4) across a chromosomal *CMV*-*GFP* transgene in HEK293 cells. Seventy-two hours after transfection with dCas9-MYB/sgRNA vectors or after treatment with Lipofectamine only, we measured GFP fluorescence via flow cytometry. Circle = median GFP fluorescence value from one transfection, 10,000 cells; bars = means of three transfections. In B and C, asterisks (*) = *p* < 0.05 for experimental mean compared to the NA control mean.

### Luciferase Assays

Luciferase assays were performed as previously described in Tekel *et al.* (37). In brief, a single well of cells from a 12 well tissue culture plate was collected per independent transfection in 1.5mL 1X PBS. Cells were loaded into 9 wells of a Black Costar Clear Bottom 96 Well Plates (Corning #3631). Three wells of cells were used to detect mCherry in order to quantify Gal4-AAP proteins. A 2X Hoechst 33342 stain (Invitrogen #H3570) was loaded into three more wells to stain nuclear DNA in order to quantify cell density. The final three wells were prepared with Luciferase Assay Buffer (Biotium #30085). Plates were scanned in a microplate reader (Biotek Synergy H1) to detect RFP (580 nm-610 nm), Hoechst 33342 fluorescence (360 nm-460 nm) and chemiluminescence from the same sample in parallel.

### RT-qPCR

We prepared total RNA from ∼1.0 × 10^6^ cells (Qiagen RNeasy Mini kit 74104) and generated cDNA from 2 µg of total RNA and the SuperScript III First Strand Synthesis system (Invitrogen #18080051) in a reaction volume of 20 μl. Quantitative PCR (qPCR) was performed with universal primers against the mCherry portion of the Gal4-AAP fusions, or the *TATA binding protein* (*TBP*) housekeeping gene. Triplicate qPCR reactions (10 μl) each contained SYBR Green 1 2X master mix (Roche), 2 µl of a 1:10 cDNA dilution, and 750 nM of each primer (forward and reverse, see Supplemental Table S5). Thermocycling conditions were as follows (minutes:seconds): pre-incubation, 95°C 05:00; amplification 45 × [98°C 00:10, 60°C 00:10, 72°C 00:10], single acquisition at 72°C; melting curve, 95°C 00:05, 65°C 01:00, 97°C continuous acquisition (5 per °C); cooling, 40°C 00:30. We calculated Mean Quantification Cycle (C_q_) for three replicate wells per unique reaction. Change in gene expression level was calculated as ΔC _q_ = 2^[Mean Cp reference − Mean Cp target]^. Log2 fold change in gene expression was calculated as = log2(ΔC _q transfected cells_ / ΔC _q untransfected_).

### Flow Cytometry

Cells were passed through a 35 μm nylon strainer (EMS #64750-25). Green fluorescent signal from GFP and red fluorescent signal from mCherry were detected on a BD Accuri C6 flow cytometer (675 nm LP filter) using CFlow Plus software. Data were further analyzed using FlowJo 10.5.3. One run (∼10 000 live cells, gated by forward and side scatter) was completed per sample, allowing us to determine median fluorescence within the live cell population.

### Construction of dCas9-MYB and Design of sgRNAs

We modified the vector pX330A_dCas9–1 × 4 (a gift from Takashi Yamamoto, Addgene plasmid #63598) by inserting a gBlock Gene Fragment (Integrated DNA Technologies) encoding the MYB TAD followed by a p2A signal (38) and *mCherry* after the *dCas9* ORF. The resulting vector expresses a dCas9-MYB fusion and mCherry as separate peptides from a single mRNA transcript. The vector and gBlock were digested with *FseI* (New England BioLabs) and FastDigest *Eco*RI (ThermoFisher Scientific) and ligated using T4 DNA Ligase (New England BioLabs). We named this new vector pX330g_dCas9-MYB. SgRNAs used in the study (Supplementary Table S3) were designed using the CRISPR design tool at crispr.mit.edu. DNA oligos were synthesized with BbsI overhangs for cloning into pX330g_dCas9-MYB (Integrative DNA Technology). Drop-in of sgRNAs followed the cloning protocol described in Cong *et al*. (39).

### Celastrol Treatments

Gal4-EED/luc cells were induced with dox and transfected as described above. Three days after transfection, cells were treated with Celastrol (Selleck Chemicals) at a final concentration of 5 μM in Gibco DMEM high glucose 1× (Life Technologies) with 10% Tet-free Fetal Bovine Serum (FBS) (Omega Scientific). Cells were incubated with the drug for six hours before being washed and either harvested for a luciferase assay or grown further in drug-free media.

### Statistical Analyses

The differences of means were calculated using the two sample, one-tailed Student’s *t* test. For *p* < 0.05, confidence was 95% for 2 degrees of freedom and a test statistic of *t*_(0.05,2)_ = 2.920. To evaluate significance of Gal4-MYB induced activation after the removal of celastrol and its subsequent re-addition, a nested one-way ANOVA was used with 95% confidence and two degrees of freedom.

## RESULTS

### Identification of Activation Associated Peptides

We surveyed public data to identify epigenetic enzymes and other proteins that are associated with transcriptional activation, and therefore might effectively disrupt repressive Polycomb chromatin. Polycomb-enriched chromatin typically includes Polycomb Repressive Complex 1 (PRC1: RING1A/B, PCGF1–PCGF6, CBX2, PHC1–PHC3, and SCMH1/2) (40), PRC2 (EZH1/2, EED, Suz12, and RBBP4/7) (40), H3K27me3, histone deacetylation, H2AK119ub1, and lncRNAs (40, 41). Each activation-associated peptide (AAP) generates modifications of histone tails either by intrinsic catalytic activity or the recruitment of chromatin-modifying co-factors. In order to predict how these AAPs might influence Polycomb heterochromatin, we searched the STRING protein-protein interaction database for binding partners and their associated chromatin-modifying activities (Supplementary Figure S1, Supplementary Table S1).

The AAPs fall into six general categories. The transcriptional activation group, (NFkB)-p65 and the MYB (c-myb) transcriptional activation domain (TAD), includes proteins that recruit RNA Polymerase II (PolII) and p300/CBP, respectively. For comparison to a strong, commonly used activator we included the recombinant TAD VP64 (four tandem copies of VP16). These AAPs have no known intrinsic gene regulation activity, and rely upon the recruitment of co-activator proteins to stimulate transcription (42–44). Histone modifications generated by the co-activators are primarily associated with a transcriptionally active state.

The histone acetylation (HAT) group includes ATF2, P300, and KAT2B. These peptides acetylate H3K27. In particular, p300 is associated with the recruitment of CBP and other co-activators that generate the activation associated mark H3K4me (45). The histone H3 methyltransferase (H3 MT) group and the H4 methyltransferase (H4 MT) group include proteins that are either Mixed-Lineage Leukemia (MLL) complex components or SET proteins. SETD7 deposits the activation associated modification H3K4me, but its regulatory impact may vary based on local DNA methylation, which can enhance or impede co-recruitment of repressive cofactors. The histone H4 methyltransferase PRMT5 induces histone acetylation that is associated with DNA methylation in some contexts (46). Still, PRMT5 primarily acts as an activator.

The final two groups, chromatin remodelers (CR) and pioneer factors (PF) represent activities that are relatively underexplored in the design of fusion-protein regulators. SMARCA4 is a chromatin remodeler that relies on an ATP-dependent reaction to shift the position of nucleosomes at a target site (47). It does not mediate the deposition of histone modifications, but is associated with CBP recruitment that evicts Polycomb-associated histone modifications (48). PFs are represented in our library by FOXA1, a winged-helix protein that displaces linker histones from DNA to facilitate a transition to open chromatin (49). In general, PFs bind to DNA within heterochromatin and do not catalyze histone post-translational modifications (28).

Several of the AAPs in our panel are associated with the eviction of Polycomb repressive complexes (PRCs) from endogenous genes. Accumulation of the chromatin remodeling protein SMARCA4 (BRG1) leads to the loss of PRCs at *Pou5f1* in mouse cells (50) and at *INK4b-ARF-INK4a* in human malignant rhabdoid tumor cells (51). In the latter case, KMT2A (MLL1) also participates in PRC depletion. ATF2 interacts with a kinase that generates H3S28p, which antagonizes PRC binding (52–54). Acetylation and methylation at H3K27 are mutually exclusive (55, 56), therefore the AAPs associated with H3K27ac (p65, MYB, ATF2, P300, KAT2B) might contribute to PRC eviction (Supplementary Figure S1). None of the AAPs in our panel are associated with enzymatic erasure of H3K27me3.

### Cis-regulatory elements recognized by transcriptional activators p65 and MYB enhance expression from an extra-chromosomal transgene

First, we used enhancer DNA elements to regulate expression from transiently-transfected plasmid DNA. Work from our group (57) and others (58, 59) has shown that plasmid DNA becomes occupied by histones, which may contribute to transgene silencing in human cells. In a previous study, we used DNA sequences that were known targets of endogenous activation-associated proteins to reduce silencing of a *luciferase* reporter gene (12). Here, we tested additional motifs (Figure 1A) that are recognized by AAPs from the transcriptional activator group in our panel: MYB and p65.

One of three MYB enhancer variants or the p65 enhancer was placed in either a forward or reverse orientation upstream of an EF1a promoter and a *luciferase* reporter (Figure 1B). PC-3 (human prostate cancer) cells were transfected with each plasmid as described previously (12). The highest mean levels of enhanced expression were observed for MYB-G-(3.4-fold, *p* = 8.6E-6), MYB-A+ (3.1-fold, *p* = 1.6E-6), MYB-C-(2.7-fold, *p* = 3.1E-4), and MYB-A-(2.3-fold, *p* = 5.0E-4) (Figure 1C). For these constructs, Luc signal values of all individual replicates were higher than the mean control value. For the remaining MYB and p65 constructs, mean Luc signal values were roughly 2-fold higher than the negative control (*p* = 9.9E-3-1.5E-2), but some of the individual replicates were at or below the mean negative control value. Overall, these results suggest that certain cis-regulatory elements from the MYB system are potent enhancers that might attract endogenous transcriptional activators to drive transgene expression from a minimal promoter.

### Identification of fusion activators with robust activity within Polycomb heterochromatin

Next, we asked whether the individual peptides MYB and p65, as well as other AAPs could enhance transgene expression in the absence of a specific enhancer sequence. To determine AAP activity within silenced chromatin, we targeted AAP fusion proteins (Supplementary Figure S2) to a chromosomal *luciferase* reporter that had been previously targeted by Polycomb repressive complexes (PRCs). The AAP open reading frames (ORFs) encode catalytic subunits or full length proteins (Supplementary Figure S2) that have been shown to support an epigenetically active state in several prior studies (42, 43, 47, 49, 61–67). All of these ORFs exclude DNA binding and histone binding domains, except for the ORF encoding FOXA1 which has a catalytic domain that requires histone interactions. We cloned each ORF into mammalian vector 14 (MV14) (Supplementary Figure S2) to express a Gal4-mCherry-AAP fusion. The Gal4 DNA binding domain serves as a module to target AAPs to UAS sequences in the transgene, while the mCherry tag allows for protein visualization and quantification of the activator fusion.

We tested all sixteen Gal4-AAP candidate fusion activators at a site that was enriched for ectopic Polycomb repressive complexes (PRCs) in HEK293 (human embryonic kidney) cells. The HEK293 cell line Gal4-EED/luc, carries a stably integrated *firefly luciferase* transgene with an upstream Gal4UAS (*Gal4UAS-Tk-luciferase*) (Figure 2A) (25, 68). The cells also carry a *TetO-CMV-Gal4EED* construct, which encodes a Gal4 DNA-binding domain (Gal4) fused to an embryonic ectoderm development (EED) open reading frame under the control of TetO-CMV promoter. The addition of doxycycline (dox) to cultured Gal4-EED/luc cells releases the TetR protein from *TetO-CMV-Gal4EED*, initiating the expression of Gal4-EED. Gal4-EED binds to the Gal4UAS site upstream of *Tk-luciferase*, recruits EZH2 (a subunit of PRC2) which generates trimethylation at histone H3 lysine 27 (H3K27me3), which in turn recruits CBX8 (a subunit of PRC2) to the reporter. Expression of *luciferase* is switched from active to silenced through accumulation of polycomb chromatin features, which have been detected by chromatin immunoprecipitation (ChIP) experiments in previous work: EZH2, Suz12, CBX8, (68), and H3K27me3 (25, 68) (Figure 2B). This well-characterized system allows us to test the activity of Gal4-AAPs with *a priori* knowledge of the chromatin environment at the target gene.

Gal4-EED/luc cells were treated with dox for two days to induce heterochromatin at the *luciferase* transgene. Afterwards, dox was removed and cells were grown for four days without dox to allow for Gal4-EED depletion. The four-day time point was chosen based on a previous report from Hansen et al. where PRC chromatin (CBX8 and H3K27me3) persisted after Gal4-EED levels had decreased (Figure 2B). Cells were then transfected with individual Gal4-AAP plasmids. *Luciferase* expression was measured three days after transfection.

Three of the sixteen Gal4-AAP-expressing samples showed increased luciferase levels compared to a untransfected control (Lipofectamine reagent only) (*p* < 0.05) (Figure 2C). To investigate whether the other Gal4 fusions were inhibited by PRC chromatin, we tested the fusion proteins at open chromatin. We used a parental HEK293 cell line, Luc14, that carries the *firefly luciferase* construct (*Gal4UAS-Tk-luciferase*) but lacks the *TetO-CMV-Gal4EED* repressor cassette (68). *Luciferase* is constitutively expressed at intermediate levels in these cells. Again, we observed that only the three Gal4-AAP fusions from the transcriptional activation group stimulated expression when targeted to the promoter-proximal UAS (Supplementary Figure S3). In both chromatin states, transcriptional activation-associated AAP’s significantly increased expression compared to the untransfected control (*p* < 0.05) by up to five-fold. Our results are consistent with other groups’ studies, where p65, VP64, or MYB stimulated gene expression from a promoter-proximal site (42–44). Here, we have demonstrated activities of these proteins within PRC-enriched chromatin.

### Gal4-MYB-induced activation at *Tk-luciferase* resists complete re-silencing over time

The results so far were obtained at a single time point after Gal4-AAP expression. We were interested in determining whether transgene activation within polycomb chromatin is stable or is transient and susceptible to eventual re-silencing (69). To investigate this question, we performed time-course experiments to measure expression from re-activated *luciferase* over time. We induced Polycomb heterochromatin in Gal4-EED/luc cells as described for the previous experiments. Two days after transfection with one of the strong activators, Gal4-VP64, -P65, or -MYB, cells were grown in dox-free medium supplemented with 10 μg/mL puromycin to select for Gal4-AAP positive cells. After three days of selection, we measured *luciferase* expression, Gal4-AAP mRNA levels, and mCherry fluorescence from a sample of each culture (Figure 3; day 3). The cells were then passaged into puromycin-free, dox-free medium to allow for the loss of Gal4-AAP, sampled every four days (approximately three cell divisions), and the same three measurements (luciferase, Gal4-AAP mRNA, and mCherry) were repeated.

For all three Gal4-AAP fusions, *luciferase* expression was significantly elevated at most time points (*p* < 0.05) compared to an untransfected control (Lipofectamine reagent only) (Figure 3A). Steep declines of Gal4-AAP mRNA and mCherry fluorescence after three days (Figure 3B and C) confirmed that the activators were transiently expressed and then depleted. Therefore, enhanced gene expression persisted to varying degrees after depletion of each Gal4-AAP, suggesting epigenetic memory of *luciferase* activation. Fluctuations in *Tk-luciferase* expression over cell culture passages suggest that the activated state is unstable after depletion of the transactivator. Four days after we ended selection for Gal4-AAP expression (day 7), Luc signal decreased roughly 3- to 4-fold, but remained significantly higher than repressed levels (control) in Gal4-VP64 and Gal4-MYB cells (Figure 3A). Luc signals spiked by roughly 2- to 4-fold at day 11 or 15, and then decreased at the next time point. We also observed fluctuations in reactivation in an additional trial (Supplementary Figure S4). This instability may be caused by competition between the activated state and background levels of Gal4-EED activity, which we have previously observed as weak levels of repression prior to dox treatment of Gal4-EED/luc cells (25). Overall, the Gal4-MYB-activated *Tk-luciferase* transgene showed the strongest resistance to re-silencing. In this case expression levels remained 1.6-fold or higher than the negative control for up to 19 days in one trial, and up to 15 days in an additional trial.

### MYB-mediated activation within closed chromatin requires interaction with a histone acetyltransferase

Next, we used a specific chemical inhibitor to probe the mechanism of MYB-driven enhancement of gene expression. The TAD core acidic domain of human MYB (D286-L309) included in our Gal4-MYB fusion construct is known to interact with a protein heterodimer of p300 and CBP (Supplementary Figure S5). A single base pair mutation within the MYB TAD domain (M303V) disrupts p300 recruitment and subsequent activation by MYB indicating that this recruitment is crucial to activation by MYB (70, 71). The p300/CBP histone acetylation complex deposits H3K27ac in opposition to H3K27me3 induced by PRC2 (72, 73). Therefore, Gal4-MYB-induced activation within Polycomb heterochromatin may be driven by histone acetylation.

To test this idea, we treated cells with a compound known to disrupt the activity of the MYB/p300/CBP complex. Celastrol is a minimally toxic pentacyclic triterpenoid that directly inhibits the MYB/p300 interaction by binding to the KIX-domain of CBP which serves as a docking site for the formation of the MYB/p300/CBP complex (74–77) (Figure 4A). Gal4-EED/luc cells were treated with dox to induce polycomb chromatin and then transfected with Gal4-MYB as described for previous experiments. We treated these cells with 5 μM celastrol for six hours. MTT assays indicated no toxicity at this concentration (Supplementary Figure S6). We observed a significant (*p* < 0.05) decrease in *luciferase* expression in celastrol-treated cells compared to an untreated control (Figure 4B). This result suggests that Gal4-MYB activity requires an interaction between MYB and p300/CBP. The other two strong activators, Gal4-VP64 and Gal4-P65, were insensitive to celastrol (Figure 4B), indicating a p300/CBP-independent mechanism for these two fusions.

In a time-course experiment we observed that Gal4-MYB activity can be switched by adding or removing celastrol from the growth medium. Eighteen hours after removal of celastrol from Gal4-MYB-transfected cells, *luciferase* expression levels increased significantly (*p* < 0.05 compared to repression at t = 6 hrs.) (Figure 4C). Re-addition of celastrol led to a reduction of Gal4-MYB-induced expression.

### MYB-mediated activation in Polycomb heterochromatin relies upon proximity to the transcriptional start site

Next we asked whether MYB-mediated activation at transgenes is context dependent. We leveraged the flexible dCas9/sgRNA system (Figure 5A) to target the MYB TAD to several sites along the *Tk-luciferase* transgene. We also tested the MYB TAD at a different transgene, *CMV-GFP* in HEK293, that had become silenced after several passages (C. Liu, unpublished).

We induced Polycomb heterochromatin in HEK 293 Gal4-EED/luc cells with dox, then removed dox to allow for Gal4-EED depletion as described above. We transfected the cells with one of four dCas9-MYB constructs, each carrying a different sgRNA targeted within the *luciferase* transgene. We measured *luciferase* expression after three days. In cells where dCas9-MYB was targeted closest to the transcription start site (+9 bp) *Tk-luciferase* expression reached the levels we observed for Gal4-MYB (Figure 5B). Expression enhancement from downstream target sites was significantly lower than Gal4-MYB (*p* < 0.05), suggesting position-dependent activity at the model Polycomb-repressed locus.

Having demonstrated dCas9-MYB activity at a PRC-repressed transgene, we set out to test MYB at endogenous heterochromatin at the *CMV-GFP* transgene. The construct, *GFP* under the control of a CMV promoter, was inserted via Cas9-mediated HDR into a non-protein-coding region of the HEK293 genome (HEK293 site 3 (78)). We transiently transfected the cells with dCas9-MYB constructs, each carrying one of four different sgRNAs targeted upstream, within the promoter, or in the coding region of the transgene. Three days after transfection, we used flow cytometry to measure GFP fluorescence compared to an untransfected control (Lipofectamine reagent only). We found that GFP fluorescence was significantly higher (*p* < 0.05) in all dCas9-MYB-expressing cells regardless of the gRNA target’s position (Figure 5C). These results indicate that MYB-mediated activation does not require proximity to the TSS in all contexts.

## DISCUSSION

We have demonstrated that DNA enhancer elements and fusion proteins derived from endogenous mammalian systems can be used to support strong expression from transiently transfected DNA. Furthermore, we have demonstrated that transient expression of Gal4-MYB confers long-term resistance to full re-silencing of a transgene in ectopic Polycomb heterochromatin. The artificial repressor (Gal4-EED) used for this study, or the incomplete erasure of certain repressive chromatin marks may have caused instability of the activated state. Future work to map chromatin features of artificially activated states over time will shed light on the requirements for stable activation. So far, our results represent some progress towards achieving reliable expression of synthetic DNA in engineered cells.

Our results also suggest that robust reactivation of a transgene within Polycomb heterochromatin is supported by the recruitment of transcription initiation complexes. However, the precise chromatin remodeling mechanism is unclear since our STRING analyses did not reveal an obvious pattern of histone modifications to distinguish the ineffective Gal4-AAPs from activators that enhanced expression in Polycomb heterochromatin (Supplementary Figure S1). Upon further investigation we determined that assembly of the MYB TAD with P300/CBP is critical for Gal4-MYB-mediated activation within Polycomb chromatin. Celastrol inhibits the interaction of p300/CBP with MYB by binding to the CBP KIX domain (74–77), and completely reduces Gal4-MYB activity. In contrast, the Gal4-P65 and Gal4-VP64 fusions showed robust activation of PRC-silenced luciferase in the presence of celastrol (Figure 5B). Although VP64 and p65 are known to interact with p300/CBP, they also interact with the large multi-subunit Mediator complex to initiate transcription (79–81). Multiple interactions of Gal4-P65 and Gal4-VP64 with Mediator may allow these proteins to function independently of p300/CBP (82). However in the case of Gal4-MYB, cooperative interactions between p300/CBP and Mediator (83, 84) may be necessary for gene activation. Mediator is known to cooperatively counteract PRC2 repression (85) and certain Mediator subunits are directly involved in the removal of PRC2 from endogenous promoters (86). Furthermore, Mediator is an antagonist of the PRC1 repression complex (87).

The inhibitor experiments also demonstrate a new technique for chemically-inducible gene regulation in mammalian cells. The ability to quickly toggle between enhanced and repressed states is a fundamental feature of engineered transgenic systems (29, 88, 89). Current methods for toggling gene expression in mammalian cells employ drug-mediated transactivator localization, such as allosteric modulation of DNA-binding protein domains (29, 88, 90), blue light-responsive CRY proteins (91), and chemically induced dimerization (CID) systems (92–94), or RNA interference to deplete the regulator (89). To our knowledge, no systems currently exist where the transactivation module’s activity (i.e., MYB-CBP binding) is modulated by a small molecule drug. Celastrol has low toxicity and is in fact being explored as a therapeutic due to its positive effects on the immune system (95–97). The concentration of celastrol that is sufficient to toggle Gal4-MYB activity in polycomb chromatin is well below the reported LD50 values for this drug (98–102).

Finally, our work demonstrates the potential flexibility of MYB fusion proteins as transactivators. dCas9-MYB showed strong activation of previously silenced transgenes near two different promoter elements, *Tk* and *CMV. Tk* had undergone silencing by ectopic polycomb chromatin, whereas *CMV* had become silenced by undetermined epigenetic factors. Interestingly, stimulation of expression from PRC-repressed *Tk* seemed to require TSS-proximal positioning of Gal4-MYB, whereas Gal4-MYB stimulated expression from both upstream (up to 1400 bp) and downstream (up to 350 bp) of the *CMV* TSS. Factors that might account for this difference include intrinsic differences in the core promoter sequences, the presence of cryptic enhancers at one promoter and not the other, and differences in chromatin structure. To our knowledge, our work represents the first use of MYB as a dCas9 fusion that can activate a transgene from proximal and distal locations.

## ACKNOWLEDGEMENTS

The authors thank C. Liu and T. Loveless for generously providing cells with a silenced CMV-GFP transgene, K. Rege and R. Niti for providing celastrol, and R. Daer and D. Vargas for early efforts on this work.

## LIST OF ABBREVIATIONS

AAP: activation associated peptide
CMV: cytomegalovirus
CR: chromatin remodeler
Gal4: Gal4 DNA binding domain
HAT: histone acetyltransferase
NLS: nuclear localization signal
ORF: open reading frame
PF: pioneer factor
PolII: RNA polymerase II
PRC: Polycomb repressive complex
TAD: transcriptional activation domain
UAS: upstream activation sequence

## Availability of data and material

Sequences of plasmids used in this study are publicly available as listed in Supplementary Table S4, and are physically available in the DNASU plasmid repository.

## Authors’ contributions

C.M.B. performed all Gal4- and dCas9-fusion cloning, HEK293 cell culture, luciferase assays, flow cytometry, and PCR related to targeted fusion activators. C.M.B. also carried out experimental design, STRING analysis, statistical analyses, and manuscript writing. J.E. designed and performed PC-3 transfections with enhancer motif-EF1a-luciferase constructs, and luciferase assays. R.M. cloned the enhancer motif-EF1a-luciferase constructs. K.A.H. oversaw all work, edited graphics for the figures, and assisted with manuscript preparation and submission. All authors reviewed and approved the manuscript.

## REFERENCES

1. Brooks, A.R., Harkins, R.N., Wang, P., Qian, H.S., Liu, P. and Rubanyi, G.M. (2004) Transcriptional silencing is associated with extensive methylation of the CMV promoter following adenoviral gene delivery to muscle. J. Gene Med., 6, 395–404.

2. Suzuki, M., Cerullo, V., Bertin, T.K., Cela, R., Clarke, C., Guenther, M., Brunetti-Pierri, N. and Lee, B. (2010) MyD88-dependent silencing of transgene expression during the innate and adaptive immune response to helper-dependent adenovirus. Hum. Gene Ther., 21, 325–336.

3. Leung, D.C. and Lorincz, M.C. (2012) Silencing of endogenous retroviruses: when and why do histone marks predominate? Trends Biochem. Sci., 37, 127–133.

4. Ross, P.J., Kennedy, M.A. and Parks, R.J. (2009) Host cell detection of noncoding stuffer DNA contained in helper-dependent adenovirus vectors leads to epigenetic repression of transgene expression. J. Virol., 83, 8409–8417.

5. Ellis, J. (2005) Silencing and variegation of gammaretrovirus and lentivirus vectors. Hum. Gene Ther., 16, 1241–1246.

6. Gong, L., Liu, F., Xiong, Z., Qi, R., Luo, Z., Gong, X., Nie, Q., Sun, Q., Liu, Y.-F., Qing, W., et al. (2018) Heterochromatin protects retinal pigment epithelium cells from oxidative damage by silencing p53 target genes. Proc. Natl. Acad. Sci. U. S. A., 115, E3987–E3995.

7. Erhardt, S., Lyko, F., Ainscough, J.F.-X., Surani, M.A. and Paro, R. (2003) Polycomb-group proteins are involved in silencing processes caused by a transgenic element from the murine imprinted H19/Igf2 region in Drosophila. Dev. Genes Evol., 213, 336–344.

8. Dufourt, J., Brasset, E., Desset, S., Pouchin, P. and Vaury, C. (2011) Polycomb group-dependent, heterochromatin protein 1-independent, chromatin structures silence retrotransposons in somatic tissues outside ovaries. DNA Res., 18, 451–461.

9. Otte, A.P. and Kwaks, T.H.J. (2003) Gene repression by Polycomb group protein complexes: a distinct complex for every occasion? Curr. Opin. Genet. Dev., 13, 448–454.

10. Johansen, J., Tornøe, J., Møller, A. and Johansen, T.E. (2003) Increased in vitro and in vivo transgene expression levels mediated through cis-acting elements. J. Gene Med., 5, 1080–1089.

11. Cheng, J.K. and Alper, H.S. (2016) Transcriptomics-Guided Design of Synthetic Promoters for a Mammalian System. ACS Synth. Biol., 5, 1455–1465.

12. Zimmerman, D., Patel, K., Hall, M. and Elmer, J. (2018) Enhancement of transgene expression by nuclear transcription factor Y and CCCTC-binding factor. Biotechnol. Prog., 10.1002/btpr.2712.

13. Wang, W., Guo, X., Li, Y.-M., Wang, X.-Y., Yang, X.-J., Wang, Y.-F. and Wang, T.-Y. (2018) Enhanced transgene expression using cis-acting elements combined with the EF1 promoter in a mammalian expression system. Eur. J. Pharm. Sci., 123, 539–545.

14. Roberts, M.L., Katsoupi, P., Tseveleki, V. and Taoufik, E. (2017) Bioinformatically Informed Design of Synthetic Mammalian Promoters. Methods Mol. Biol., 1651, 93–112.

15. Saxena, P., Bojar, D. and Fussenegger, M. (2017) Design of Synthetic Promoters for Gene Circuits in Mammalian Cells. In Methods in Molecular Biology .pp. 263–273.

16. Morita, S., Noguchi, H., Horii, T., Nakabayashi, K., Kimura, M., Okamura, K., Sakai, A., Nakashima, H., Hata, K., Nakashima, K., et al. (2016) Targeted DNA demethylation in vivo using dCas9-peptide repeat and scFv-TET1 catalytic domain fusions. Nat. Biotechnol., 34, 1060–1065.

17. Liu, P., Chen, M., Liu, Y., Qi, L.S. and Ding, S. (2018) CRISPR-Based Chromatin Remodeling of the Endogenous Oct4 or Sox2 Locus Enables Reprogramming to Pluripotency. Cell Stem Cell, 22, 252–261.e4.

18. Hilton, I.B., D’Ippolito, A.M., Vockley, C.M., Thakore, P.I., Crawford, G.E., Reddy, T.E. and Gersbach, C.A. (2015) Epigenome editing by a CRISPR-Cas9-based acetyltransferase activates genes from promoters and enhancers. Nat. Biotechnol., 33, 510–517.

19. Chavez, A., Scheiman, J., Vora, S., Pruitt, B.W., Tuttle, M., Iyer, E., Kiani, S., Guzman, C.D., Wiegand, D.J., Ter-Ovanesyan, D., et al. (2014) Highly-efficient Cas9-mediated transcriptional programming. 10.1101/012880.

20. Konermann, S., Brigham, M.D., Trevino, A.E., Joung, J., Abudayyeh, O.O., Barcena, C., Hsu, P.D., Habib, N., Gootenberg, J.S., Nishimasu, H., et al. (2015) Genome-scale transcriptional activation by an engineered CRISPR-Cas9 complex. Nature, 517, 583–588.

21. Huang, Y.-H., Su, J., Lei, Y., Brunetti, L., Gundry, M.C., Zhang, X., Jeong, M., Li, W. and Goodell, M.A. (2017) DNA epigenome editing using CRISPR-Cas SunTag-directed DNMT3A. Genome Biol., 18, 176.

22. Black, J.B., Adler, A.F., Wang, H.-G., D’Ippolito, A.M., Hutchinson, H.A., Reddy, T.E., Pitt, G.S., Leong, K.W. and Gersbach, C.A. (2016) Targeted Epigenetic Remodeling of Endogenous Loci by CRISPR/Cas9-Based Transcriptional Activators Directly Converts Fibroblasts to Neuronal Cells. Cell Stem Cell, 19, 406–414.

23. Gao, X., Tsang, J.C.H., Gaba, F., Wu, D., Lu, L. and Liu, P. (2014) Comparison of TALE designer transcription factors and the CRISPR/dCas9 in regulation of gene expression by targeting enhancers. Nucleic Acids Res., 42, e155.

24. Zhang, Y., Yin, C., Zhang, T., Li, F., Yang, W., Kaminski, R., Fagan, P.R., Putatunda, R., Young, W.-B., Khalili, K., et al. (2015) CRISPR/gRNA-directed synergistic activation mediator (SAM) induces specific, persistent and robust reactivation of the HIV-1 latent reservoirs. Sci. Rep., 5, 16277.

25. Daer, R.M., Cutts, J.P., Brafman, D.A. and Haynes, K.A. (2017) The Impact of Chromatin Dynamics on Cas9-Mediated Genome Editing in Human Cells. ACS Synth. Biol., 6, 428–438.

26. Clapier, C.R., Iwasa, J., Cairns, B.R. and Peterson, C.L. (2017) Mechanisms of action and regulation of ATP-dependent chromatin-remodelling complexes. Nat. Rev. Mol. Cell Biol., 18, 407–422.

27. Zaret, K.S. and Carroll, J.S. (2011) Pioneer transcription factors: establishing competence for gene expression. Genes Dev., 25, 2227–2241.

28. Magnani, L., Eeckhoute, J. and Lupien, M. (2011) Pioneer factors: directing transcriptional regulators within the chromatin environment. Trends Genet., 27, 465–474.

29. Kramer, B.P., Viretta, A.U., Daoud-El-Baba, M., Aubel, D., Weber, W. and Fussenegger, M. (2004) An engineered epigenetic transgene switch in mammalian cells. Nat. Biotechnol., 22, 867–870.

30. Kwaks, T.H.J., Sewalt, R.G.A.B., van Blokland, R., Siersma, T.J., Kasiem, M., Kelder, A. and Otte, A.P. (2005) Targeting of a histone acetyltransferase domain to a promoter enhances protein expression levels in mammalian cells. J. Biotechnol., 115, 35–46.

31. Santillan, D.A., Theisler, C.M., Ryan, A.S., Popovic, R., Stuart, T., Zhou, M.-M., Alkan, S. and Zeleznik-Le, N.J. (2006) Bromodomain and histone acetyltransferase domain specificities control mixed lineage leukemia phenotype. Cancer Res., 66, 10032–10039.

32. Cano-Rodriguez, D., Gjaltema, R.A.F., Jilderda, L.J., Jellema, P., Dokter-Fokkens, J., Ruiters, M.H.J. and Rots, M.G. (2016) Writing of H3K4Me3 overcomes epigenetic silencing in a sustained but context-dependent manner. Nat. Commun., 7.

33. McGinty, R.K., Kim, J., Chatterjee, C., Roeder, R.G. and Muir, T.W. (2008) Chemically ubiquitylated histone H2B stimulates hDot1L-mediated intranucleosomal methylation. Nature, 453, 812–816.

34. O’Geen, H., Ren, C., Nicolet, C.M., Perez, A.A., Halmai, J., Le, V.M., Mackay, J.P., Farnham, P.J. and Segal, D.J. (2017) dCas9-based epigenome editing suggests acquisition of histone methylation is not sufficient for target gene repression. Nucleic Acids Res., 45, 9901–9916.

35. Aloia, L., Di Stefano, B. and Di Croce, L. (2013) Polycomb complexes in stem cells and embryonic development. Development, 140, 2525–2534.

36. Poynter, S.T. and Kadoch, C. (2016) Polycomb and trithorax opposition in development and disease. Wiley Interdiscip. Rev. Dev. Biol., 5, 659–688.

37. Tekel, S.J., Barrett, C., Vargas, D. and Haynes, K.A. (2018) Design, Construction, and Validation of Histone-Binding Effectors in Vitro and in Cells. Biochemistry, 57, 4707–4716.

38. Liu, Z., Chen, O., Blake Joseph Wall, J., Zheng, M., Zhou, Y., Wang, L., Vaseghi, H.R., Qian, L. and Liu, J. (2017) Systematic comparison of 2A peptides for cloning multi-genes in a polycistronic vector. Sci. Rep., 7.

39. Cong, L., Ran, F.A., Cox, D., Lin, S., Barretto, R., Habib, N., Hsu, P.D., Wu, X., Jiang, W., Marraffini, L.A., et al. (2013) Multiplex genome engineering using CRISPR/Cas systems. Science, 339, 819–823.

40. Schuettengruber, B., Bourbon, H.-M., Di Croce, L. and Cavalli, G. (2017) Genome Regulation by Polycomb and Trithorax: 70 Years and Counting. Cell, 171, 34–57.

41. Simon, J.A. and Kingston, R.E. (2013) Occupying chromatin: Polycomb mechanisms for getting to genomic targets, stopping transcriptional traffic, and staying put. Mol. Cell, 49, 808–824.

42. Beerli, R.R., Segal, D.J., Dreier, B. and Barbas, C.F., 3rd (1998) Toward controlling gene expression at will: specific regulation of the erbB-2/HER-2 promoter by using polydactyl zinc finger proteins constructed from modular building blocks. Proc. Natl. Acad. Sci. U. S. A., 95, 14628–14633.

43. Liu, P.Q., Rebar, E.J., Zhang, L., Liu, Q., Jamieson, A.C., Liang, Y., Qi, H., Li, P.X., Chen, B., Mendel, M.C., et al. (2001) Regulation of an endogenous locus using a panel of designed zinc finger proteins targeted to accessible chromatin regions. Activation of vascular endothelial growth factor A. J. Biol. Chem., 276, 11323–11334.

44. Weston, K. and Michael Bishop, J. (1989) Transcriptional activation by the v-myb oncogene and its cellular progenitor, c-myb. Cell, 58, 85–93.

45. Vo, N. and Goodman, R.H. (2001) CREB-binding Protein and p300 in Transcriptional Regulation. J. Biol. Chem., 276, 13505–13508.

46. Zhao, Q., Rank, G., Tan, Y.T., Li, H., Moritz, R.L., Simpson, R.J., Cerruti, L., Curtis, D.J., Patel, D.J., Allis, C.D., et al. (2009) PRMT5-mediated methylation of histone H4R3 recruits DNMT3A, coupling histone and DNA methylation in gene silencing. Nat. Struct. Mol. Biol., 16, 304–311.

47. Antonysamy, S., Bonday, Z., Campbell, R.M., Doyle, B., Druzina, Z., Gheyi, T., Han, B., Jungheim, L.N., Qian, Y., Rauch, C., et al. (2012) Crystal structure of the human PRMT5:MEP50 complex. Proc. Natl. Acad. Sci. U. S. A., 109, 17960–17965.

48. Alver, B.H., Kim, K.H., Lu, P., Wang, X., Manchester, H.E., Wang, W., Haswell, J.R., Park, P.J. and Roberts, C.W.M. (2017) The SWI/SNF chromatin remodelling complex is required for maintenance of lineage specific enhancers. Nat. Commun., 8, 14648.

49. Clark, K.L., Halay, E.D., Lai, E. and Burley, S.K. (1993) Co-crystal structure of the HNF-3/fork head DNA-recognition motif resembles histone H5. Nature, 364, 412–420.

50. Kadoch, C., Williams, R.T., Calarco, J.P., Miller, E.L., Weber, C.M., Braun, S.M.G., Pulice, J.L., Chory, E.J. and Crabtree, G.R. (2017) Dynamics of BAF-Polycomb complex opposition on heterochromatin in normal and oncogenic states. Nat. Genet., 49, 213–222.

51. Kia, S.K., Gorski, M.M., Giannakopoulos, S. and Verrijzer, C.P. (2008) SWI/SNF mediates polycomb eviction and epigenetic reprogramming of the INK4b-ARF-INK4a locus. Mol. Cell. Biol., 28, 3457–3464.

52. Lau, P.N.I. and Cheung, P. (2011) Histone code pathway involving H3 S28 phosphorylation and K27 acetylation activates transcription and antagonizes polycomb silencing. Proc. Natl. Acad. Sci. U. S. A., 108, 2801–2806.

53. Gehani, S.S., Agrawal-Singh, S., Dietrich, N., Christophersen, N.S., Helin, K. and Hansen, K. (2010) Polycomb group protein displacement and gene activation through MSK-dependent H3K27me3S28 phosphorylation. Mol. Cell, 39, 886–900.

54. Josefowicz, S.Z., Shimada, M., Armache, A., Li, C.H., Miller, R.M., Lin, S., Yang, A., Dill, B.D., Molina, H., Park, H.-S., et al. (2016) Chromatin Kinases Act on Transcription Factors and Histone Tails in Regulation of Inducible Transcription. Mol. Cell, 64, 347–361.

55. Tie, F., Banerjee, R., Stratton, C.A., Prasad-Sinha, J., Stepanik, V., Zlobin, A., Diaz, M.O., Scacheri, P.C. and Harte, P.J. (2009) CBP-mediated acetylation of histone H3 lysine 27 antagonizes Drosophila Polycomb silencing. Development, 136, 3131–3141.

56. Englert, N.A., Luo, G., Goldstein, J.A. and Surapureddi, S. (2014) Epigenetic Modification of Histone 3 Lysine 27. J. Biol. Chem., 290, 2264–2278.

57. Christensen, M.D., Nitiyanandan, R., Meraji, S., Daer, R., Godeshala, S., Goklany, S., Haynes, K. and Rege, K. (2018) An inhibitor screen identifies histone-modifying enzymes as mediators of polymer-mediated transgene expression from plasmid DNA. J. Control. Release, 286, 210–223.

58. Gracey Maniar, L.E., Maniar, J.M., Chen, Z.-Y., Lu, J., Fire, A.Z. and Kay, M.A. (2013) Minicircle DNA vectors achieve sustained expression reflected by active chromatin and transcriptional level. Mol. Ther., 21, 131–138.

59. Riu, E., Chen, Z.-Y., Xu, H., He, C.-Y. and Kay, M.A. (2007) Histone modifications are associated with the persistence or silencing of vector-mediated transgene expression in vivo. Mol. Ther., 15, 1348–1355.

60. Mathelier, A., Fornes, O., Arenillas, D.J., Chen, C.-Y., Denay, G., Lee, J., Shi, W., Shyr, C., Tan, G., Worsley-Hunt, R., et al. (2016) JASPAR 2016: a major expansion and update of the open–access database of transcription factor binding profiles. Nucleic Acids Res., 44, D110–5.

61. Zobel, A., Kalkbrenner, F., Vorbrueggen, G. and Moelling, K. (1992) Transactivation of the human c-myc gene by c-Myb. Biochem. Biophys. Res. Commun., 186, 715–722.

62. Kawasaki, H., Schiltz, L., Chiu, R., Itakura, K., Taira, K., Nakatani, Y. and Yokoyama, K.K. (2000) ATF-2 has intrinsic histone acetyltransferase activity which is modulated by phosphorylation. Nature, 405, 195–200.

63. Yang, L., Wang, H., Luo, X., Mao, P., Tian, W., Shi, Y., Huang, G., Zhang, J. and Ma, D. (2011) Virion protein 16 induces demethylation of DNA integrated within chromatin in a novel mammalian cell model. Acta Biochim. Biophys. Sin., 44, 154–161.

64. Milne, T.A., Briggs, S.D., Brock, H.W., Martin, M.E., Gibbs, D., Allis, C.D. and Hess, J.L. (2002) MLL targets SET domain methyltransferase activity to Hox gene promoters. Mol. Cell, 10, 1107–1117.

65. Nishioka, K., Chuikov, S., Sarma, K., Erdjument-Bromage, H., Allis, C.D., Tempst, P. and Reinberg, D. (2002) Set9, a novel histone H3 methyltransferase that facilitates transcription by precluding histone tail modifications required for heterochromatin formation. Genes Dev., 16, 479–489.

66. Serandour, A.A., Avner, S., Percevault, F., Demay, F., Bizot, M., Lucchetti-Miganeh, C., Barloy-Hubler, F., Brown, M., Lupien, M., Metivier, R., et al. (2011) Epigenetic switch involved in activation of pioneer factor FOXA1-dependent enhancers. Genome Res., 21, 555–565.

67. Wang, W., Côté, J., Xue, Y., Zhou, S., Khavari, P.A., Biggar, S.R., Muchardt, C., Kalpana, G.V., Goff, S.P., Yaniv, M., et al. (1996) Purification and biochemical heterogeneity of the mammalian SWI-SNF complex. EMBO J., 15, 5370–5382.

68. Hansen, K.H., Bracken, A.P., Pasini, D., Dietrich, N., Gehani, S.S., Monrad, A., Rappsilber, J., Lerdrup, M. and Helin, K. (2008) A model for transmission of the H3K27me3 epigenetic mark. Nat. Cell Biol., 10, 1291–1300.

69. Pirrotta, V. (2017) Introduction to Polycomb Group Mechanisms. In Polycomb Group Proteins .pp. 1–3.

70. Sandberg, M.L., Sutton, S.E., Pletcher, M.T., Wiltshire, T., Tarantino, L.M., Hogenesch, J.B. and Cooke, M.P. (2005) c-Myb and p300 regulate hematopoietic stem cell proliferation and differentiation. Dev. Cell, 8, 153–166.

71. Pattabiraman, D.R., Sun, J., Dowhan, D.H., Ishii, S. and Gonda, T.J. (2009) Mutations in multiple domains of c-Myb disrupt interaction with CBP/p300 and abrogate myeloid transforming ability. Mol. Cancer Res., 7, 1477–1486.

72. Raisner, R., Kharbanda, S., Jin, L., Jeng, E., Chan, E., Merchant, M., Haverty, P.M., Bainer, R., Cheung, T., Arnott, D., et al. (2018) Enhancer Activity Requires CBP/P300 Bromodomain-Dependent Histone H3K27 Acetylation. Cell Rep., 24, 1722–1729.

73. Ogryzko, V.V., Louis Schiltz, R., Russanova, V., Howard, B.H. and Nakatani, Y. (1996) The Transcriptional Coactivators p300 and CBP Are Histone Acetyltransferases. Cell, 87, 953–959.

74. Coulibaly, A., Haas, A., Steinmann, S., Jakobs, A., Schmidt, T.J. and Klempnauer, K.-H. (2018) The natural anti-tumor compound Celastrol targets a Myb-C/EBPβ-p300 transcriptional module implicated in myeloid gene expression. PLoS One, 13, e0190934.

75. Uttarkar, S., Piontek, T., Dukare, S., Schomburg, C., Schlenke, P., Berdel, W.E., Müller-Tidow, C., Schmidt, T.J. and Klempnauer, K.-H. (2016) Small-Molecule Disruption of the Myb/p300 Cooperation Targets Acute Myeloid Leukemia Cells. Mol. Cancer Ther., 15, 2905–2915.

76. Denis, C.M., Langelaan, D.N., Kirlin, A.C., Chitayat, S., Munro, K., Spencer, H.L., LeBrun, D.P. and Smith, S.P. (2014) Functional redundancy between the transcriptional activation domains of E2A is mediated by binding to the KIX domain of CBP/p300. Nucleic Acids Res., 42, 7370–7382.

77. Uttarkar, S., Dassé, E., Coulibaly, A., Steinmann, S., Jakobs, A., Schomburg, C., Trentmann, A., Jose, J., Schlenke, P., Berdel, W.E., et al. (2016) Targeting acute myeloid leukemia with a small molecule inhibitor of the Myb/p300 interaction. Blood, 127, 1173–1182.

78. Tsai, S.Q., Zheng, Z., Nguyen, N.T., Liebers, M., Topkar, V.V., Thapar, V., Wyvekens, N., Khayter, C., Iafrate, A.J., Le, L.P., et al. (2015) GUIDE-seq enables genome-wide profiling of off-target cleavage by CRISPR-Cas nucleases. Nat. Biotechnol., 33, 187–197.

79. van Essen, D., Engist, B., Natoli, G. and Saccani, S. (2009) Two Modes of Transcriptional Activation at Native Promoters by NF-κB p65. PLoS Biol., 7, e1000073.

80. Thakore, P.I., Black, J.B., Hilton, I.B. and Gersbach, C.A. (2016) Editing the epigenome: technologies for programmable transcription and epigenetic modulation. Nat. Methods, 13, 127–137.

81. Wang, W., Qin, J.-J., Voruganti, S., Nag, S., Zhou, J. and Zhang, R. (2015) Polycomb Group (PcG) Proteins and Human Cancers: Multifaceted Functions and Therapeutic Implications. Med. Res. Rev., 35, 1220–1267.

82. Lecoq, L., Raiola, L., Chabot, P.R., Cyr, N., Arseneault, G., Legault, P. and Omichinski, J.G. (2017) Structural characterization of interactions between transactivation domain 1 of the p65 subunit of NF-κB and transcription regulatory factors. Nucleic Acids Res., 45, 5564–5576.

83. Huang, Z.-Q., Li, J., Sachs, L.M., Cole, P.A. and Wong, J. (2003) A role for cofactor-cofactor and cofactor-histone interactions in targeting p300, SWI/SNF and Mediator for transcription. EMBO J., 22, 2146–2155.

84. Haas, M., Siegert, M., Schürmann, A., Sodeik, B. and Wolfes, H. (2004) c-Myb protein interacts with Rcd-1, a component of the CCR4 transcription mediator complex. Biochemistry, 43, 8152–8159.

85. Fukasawa, R., Iida, S., Tsutsui, T., Hirose, Y. and Ohkuma, Y. (2015) Mediator complex cooperatively regulates transcription of retinoic acid target genes with Polycomb Repressive Complex 2 during neuronal differentiation. J. Biochem., 158, 373–384.

86. Englert, N.A., Luo, G., Goldstein, J.A. and Surapureddi, S. (2015) Epigenetic modification of histone 3 lysine 27: mediator subunit MED25 is required for the dissociation of polycomb repressive complex 2 from the promoter of cytochrome P450 2C9. J. Biol. Chem., 290, 2264–2278.

87. Lehmann, L., Ferrari, R., Vashisht, A.A., Wohlschlegel, J.A., Kurdistani, S.K. and Carey, M. (2012) Polycomb repressive complex 1 (PRC1) disassembles RNA polymerase II preinitiation complexes. J. Biol. Chem., 287, 35784–35794.

88. Oakes, B.L., Nadler, D.C., Flamholz, A., Fellmann, C., Staahl, B.T., Doudna, J.A. and Savage, D.F. (2016) Profiling of engineering hotspots identifies an allosteric CRISPR-Cas9 switch. Nat. Biotechnol., 34, 646–651.

89. Greber, D., El-Baba, M.D. and Fussenegger, M. (2008) Intronically encoded siRNAs improve dynamic range of mammalian gene regulation systems and toggle switch. Nucleic Acids Res., 36, e101–e101.

90. Stanton, B.C., Siciliano, V., Ghodasara, A., Wroblewska, L., Clancy, K., Trefzer, A.C., Chesnut, J.D., Weiss, R. and Voigt, C.A. (2014) Systematic transfer of prokaryotic sensors and circuits to mammalian cells. ACS Synth. Biol., 3, 880–891.

91. Mansouri, M., Strittmatter, T. and Fussenegger, M. (2018) Light-Controlled Mammalian Cells and Their Therapeutic Applications in Synthetic Biology. Adv. Sci. Lett.

92. Inobe, T. and Nukina, N. (2016) Rapamycin-induced oligomer formation system of FRB-FKBP fusion proteins. J. Biosci. Bioeng., 122, 40–46.

93. Rivera, V.M., Clackson, T., Natesan, S., Pollock, R., Amara, J.F., Keenan, T., Magari, S.R., Phillips, T., Courage, N.L., Cerasoli, F., et al. (1996) A humanized system for pharmacologic control of gene expression. Nat. Med., 2, 1028–1032.

94. DeRose, R., Miyamoto, T. and Inoue, T. (2013) Manipulating signaling at will: chemically-inducible dimerization (CID) techniques resolve problems in cell biology. Pflugers Arch., 465, 409–417.

95. Cascão, R., Fonseca, J.E. and Moita, L.F. (2017) Celastrol: A Spectrum of Treatment Opportunities in Chronic Diseases. Front. Med., 4, 69.

96. Venkatesha, S.H., Dudics, S., Astry, B. and Moudgil, K.D. (2016) Control of autoimmune inflammation by celastrol, a natural triterpenoid. Pathog. Dis., 74.

97. Ju, S.M., Youn, G.S., Cho, Y.S., Choi, S.Y. and Park, J. (2015) Celastrol ameliorates cytokine toxicity and pro-inflammatory immune responses by suppressing NF-κB activation in RINm5F beta cells. BMB Rep., 48, 172–177.

98. Li, G., Liu, D., Zhang, Y., Qian, Y., Zhang, H., Guo, S., Sunagawa, M., Hisamitsu, T. and Liu, Y. (2013) Celastrol inhibits lipopolysaccharide-stimulated rheumatoid fibroblast-like synoviocyte invasion through suppression of TLR4/NF-κB-mediated matrix metalloproteinase-9 expression. PLoS One, 8, e68905.

99. Raja, S.M., Clubb, R.J., Ortega-Cava, C., Williams, S.H., Bailey, T.A., Duan, L., Zhao, X., Reddi, A.L., Nyong, A.M., Natarajan, A., et al. (2011) Anticancer activity of Celastrol in combination with ErbB2-targeted therapeutics for treatment of ErbB2-overexpressing breast cancers. Cancer Biol. Ther., 11, 263–276.

100. Yang, H., Chen, D., Cui, Q.C., Yuan, X. and Dou, Q.P. (2006) Celastrol, a triterpene extracted from the Chinese ‘Thunder of God Vine,’ is a potent proteasome inhibitor and suppresses human prostate cancer growth in nude mice. Cancer Res., 66, 4758–4765.

101. Cleren, C., Calingasan, N.Y., Chen, J. and Beal, M.F. (2005) Celastrol protects against MPTP- and 3-nitropropionic acid-induced neurotoxicity. J. Neurochem., 94, 995–1004.

102. Konieczny, J., Jantas, D., Lenda, T., Domin, H., Czarnecka, A., Kuter, K., Smialowska, M., Lasoń, W. and Lorenc-Koci, E. (2014) Lack of neuroprotective effect of celastrol under conditions of proteasome inhibition by lactacystin in in vitro and in vivo studies: implications for Parkinson’s disease. Neurotox. Res., 26, 255–273.

