## Supplementary Tables S2 - S5 and Figures S1 - S5 for "Components from the human c-myb transcriptional regulation system reactivate epigenetically repressed transgenes"

**SUPPLEMENTARY INFORMATION**

Table S2. Activation-associated peptide (AAP) accession numbers and primers

Table S3. dCas9-MYB sgRNA targeting sequences

Table S4. Publicly accessible plasmids used in this study

Table S5. Primers for qPCR to quantify Gal4-AAP mRNA

Figure S1. Histone modifications associated with activation-associated peptides (AAPs) used in this study

Figure S2. Design and construction of Activation-associated peptide (AAP) -Gal4 fusions

Figure S3. Measurement of luciferase reporter expression within open chromatin after exposure to Gal4-AAP fusions.

Figure S3. Additional trial of time course experiments with Gal4-AAP-expressing cells

Figure S4. Annotated motifs within the MYB protein

Figure S5. MTT cell viability assays

**Table S2**. Activation-associated peptide (AAP) Accession Numbers and Primers. Capitalized nucleotides indicate overhangs for the addition of XbaI and NotI sites for cloning.

| **AAP** | **Ref.** | **UniProt ID** | **Isoform** | **NCBI RefSeq** | **Amplification primers (5’ .. 3’)** |
| --- | --- | --- | --- | --- | --- |
| VP64 (4xVP16) | [S1] | n.a. | - | - | F: CCTTATCTAGAgacgctttggacgacttcga  R: CTTAGCGGCCGacaacatgtccaagtcgaagt |
| (NKκB)-p65 | [S2] | Q04206 | - | NM_001145138.1 | F: CCTTATCTAGAtacctgccagatacagacga  R: CCTTAGCGGCCGatctcagccctgctgagtcag |
| MYB | [S3] | P10242 | Isoform 1 P10242-1 | NM_005375.2 | F: GGCCTTATCTAGccagctgccgcagccattca  R: AATTAGCGGCCGcccacccggggtagctgcat |
| ATF2 | [S4] | P15336 | Isoform 1 P15336-1 | NM_001256090.1 | F: CCTTATCTAGAgagatgacactgaaatttgg  R: CCTTAGCGGCCGaggatcttcgttagctgctc |
| p300 | [S4] | Q09472 | Q09472-1 | NM_001429.3 | gBlock from IDT |
| KAT2B (PCAF) | [S5] | Q92831 | Q92831-1 | NM_003884.4 | F: CCTTATCTAGActcaaccagaaaccaaacaa  R: CCTTAGCGGCCGatttagctcacatcccatta |
| KMT2A (MLL1) | [S6] | Q03164 | Isoform 1 Q03164-1 | NM_005933.3 | gBlock from IDT |
| KMT2C (MML3) | [S6] | Q8NEZ4 | Isoform 1 Q8NEZ4-1 | NM_170606.2 | gBlock from IDT |
| KMT2D (MLL2, 4) | [S6] | O14686 | Isoform 1 O14686-1 | NM_003482.3 | gBlock from IDT |
| KMT2E (MLL5) | [S6] | Q8IZD2 | Isoform 1 Q8IZD2-1 | NM_182931.2 (variant 1) | F: GCGCTCTAGAaatttggataaagagagggc  R: AATGCGGCCGcttcattactaataggagtt |
| SETD1A (SET1) | [S6] | O15047 | O15047-1 | NM_014712.1 | F: CCTTATCTAGAaagaagctccgatttggccg  R: CCTTAGCGGCCGagtttagggagccccggcagc |
| SETD1B (SET1B) | [S6] | Q9UPS6 | Isoform 1 Q9UPS6-1 | No RefSeq number | gBlock from IDT |
| SETD7 (SET7/9) | [S7] | Q8WTS6 | Q8WTS6-1 | NM_030648.2 | gBlock from IDT |
| PRMT5 | [S8] | O14744 | Isoform 1 O14744-1 | NM_006109.4 | F: ATGCTCTAGAatggcggcgatggcggtcgg  R: ATGCGCGGCCGcgaggccaatggtatatgagc |
| FOXA1 | [S9, S10] | P55317 | P55317-1 | NM_004496 | F: GGCTCTAGAatgttaggaactgtgaagatgga  R: ATTGCGGCCGccaaggaagtgtttaggacgg |
| SMARCA4 (BRG1) | [S11] | P51532 | Isoform 1 P51532-1 | NM_001128844.1 (variant 2) | F: CCTTATCTAGAatgtccactccagacccacc  R: CCTTAGCGGCCGgctgctgtccttgtacttga |

**Table S3**. dCas9-MYB sgRNA targeting sequences. The targeting sequences for g46, g31, g32, and g25 have been used in our previous work [S12].

| **Transgene** | **Cell line** | **Target Name** | **Targeting Sequence** |
| --- | --- | --- | --- |
| *Tk-Luciferase* | HEK293 Gal4-EED/luc [S13] | g46 | 5’ cctgcataagcttgccacca 3’ |
|  |  | g31 | 5’ cgaggtgaacatcacgtacg 3’ |
|  |  | g32 | 5’ cctctagaggatggaaccgc 3’ |
|  |  | g25 | 5’ accgtagtgtttgtttccaa 3’ |
| *CMV-GFP* | HEK293 GFP (Chang Liu, UC Irvine, unpublished) | L1 | 5’ ctctgtcacaggactcagcc 3’ |
|  |  | L2 | 5’ gggcggtaggcgtgtacggt 3’ |
|  |  | L3 | 5’ cactggtgtcgtccctattc 3’ |
|  |  | L4 | 5’ cttcttcaagtccgccatgc 3’ |

**Table S4**. Publicly accessible plasmids. Annotated sequences can be viewed online in the Haynes lab Benchling collection “DNA-Binding Fusion Transcriptional Regulators” at <https://benchling.com/hayneslab/f_/5wovkOaK-gal4-dna-binding-fusion-transcription-regulators/?sort=name>.

| **Name** | **DNASU Accession #** |
| --- | --- |
| MV14 | - |
| MV14_VP64 | HaCD00812388 |
| MV14_p65 | HsCD00812387 |
| MV14_MYB | - |
| MV14_ATF2 | HsCD00833013 |
| MV14_p300 | HsCD00833014 |
| MV14_KAT2B | HsCD00833015 |
| MV14_KMT2A | HsCD00833016 |
| MV14_KMT2C | HsCD00833018 |
| MV14_KMT2D | HsCD00833017 |
| MV14_KMT2E | - |
| MV14_SETD1A | HsCD00833019 |
| MV14_SETD1B | HsCD00833020 |
| MV14_SETD7 | HsCD00833021 |
| MV14_PRMT5 | - |
| MV14_FOXA1 | - |
| MV14_SMARCA4 | HsCD00833022 |
| pX330g_dCas9_MYB | - |

**Table S5.** Primers for RT-qPCR

| **Target** | **Amplification primers (5’ .. 3’)** |
| --- | --- |
| *mCherry* (Gal4-AAP transcripts) | F: gctccaaggcctacgtgaag  R: aagttcatcacgcgctccca |
| *TBP* (housekeeping gene) | F: caggggttcagtgaggtcg  R: ccctgggtcactgcaaagat |

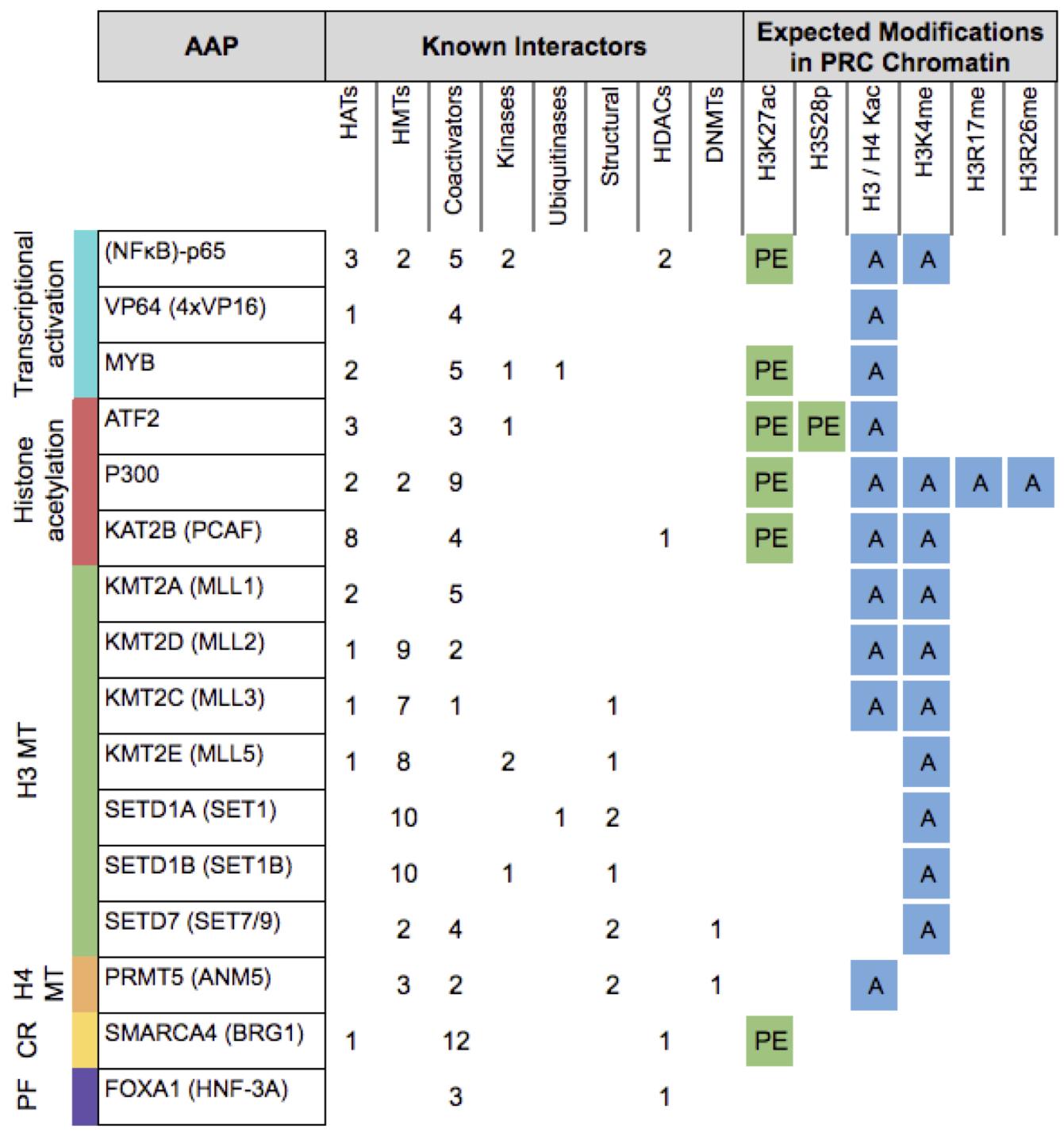

**Figure S1.** Histone modifications associated with activation-associated peptides (AAPs) used in this study. Interaction partners determined by STRING analysis are listed in Supplementary Table S1. Previous work that characterized each AAP is cited in Supplementary Table S2. H3 MT = histone H3 methyltransferase, H4 MT = histone H4 methyltransferase, CR = chromatin remodeler, PF = pioneer factor, PE = Polycomb eviction, A = transcriptional activation.

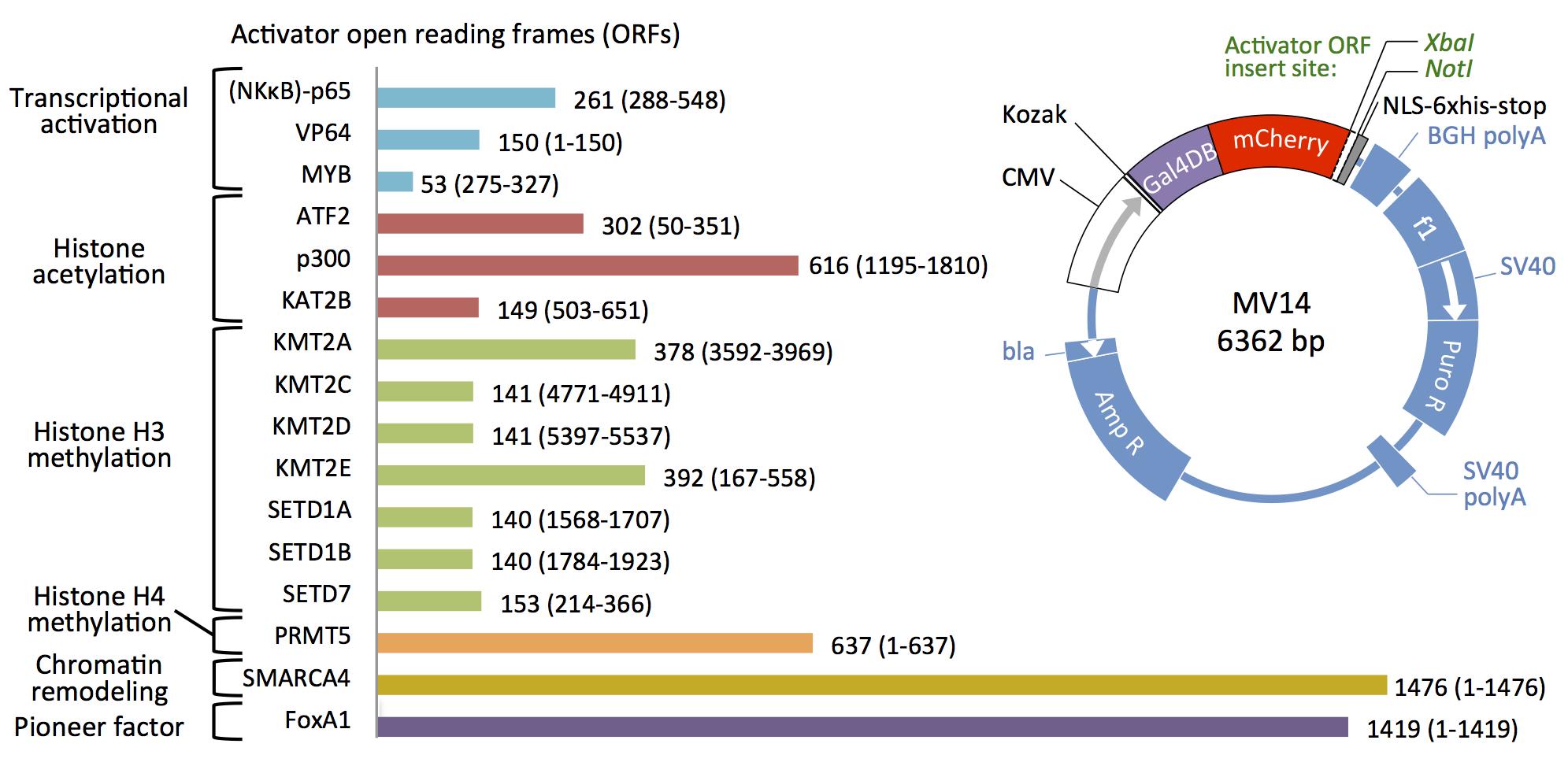

**Figure S2.** Design and construction of Activation-associated peptide (AAP) -Gal4 fusions. Amino acid lengths are indicated by the lengths of the colored bars and the number at the end of each bar. The AAP domain within the full length wild-type sequence (Supplementary Table S2) is indicated by amino acid positions in parenthesis. ORFs were cloned into MV14 to express a Gal4-AAP fusion protein from a cytomegalovirus (CMV) promoter. Gal4-AAPs are expressed with a C-terminal nuclear localization signal (NLS) and a 6X histidine tag. MV14 expresses puromycin resistance to enable selection of Gal4-AAP positive cells.

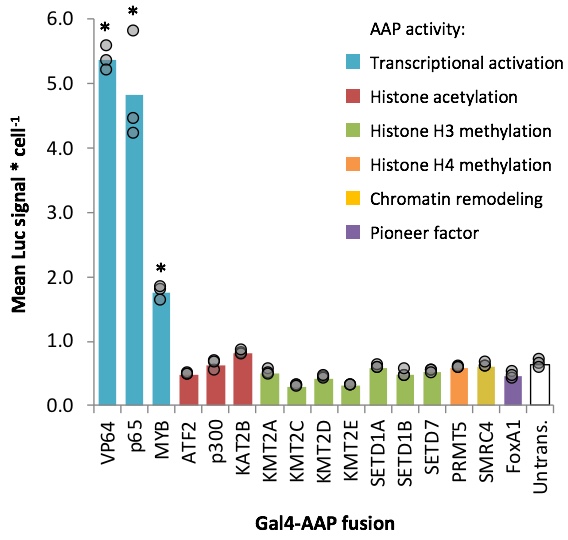

**Figure S3**. Measurement of luciferase reporter expression within open chromatin after exposure to Gal4-AAP fusions. Luc14 cells carry the same *Tk-luciferase* locus at the same chromosome site as Gal4-EED/luc cells, but without Gal4-EED. Therefore, the site is PRC-depleted and basal levels of expression from *Tk-luciferase* are higher than in Gal4-EED/luc cells (Figure 2). Luc14 cells were transfected with each Gal4-AAP fusion plasmid. Three days after transfection luciferase signal was measured. Each circle in the bar graph shows the mean luciferase (Luc) signal for a single transfection, divided by cell density (total DNA, Hoechst staining signal). Bars show means of three transfections. Asterisks (*) = *p* < 0.05 compared to untransfected cells.

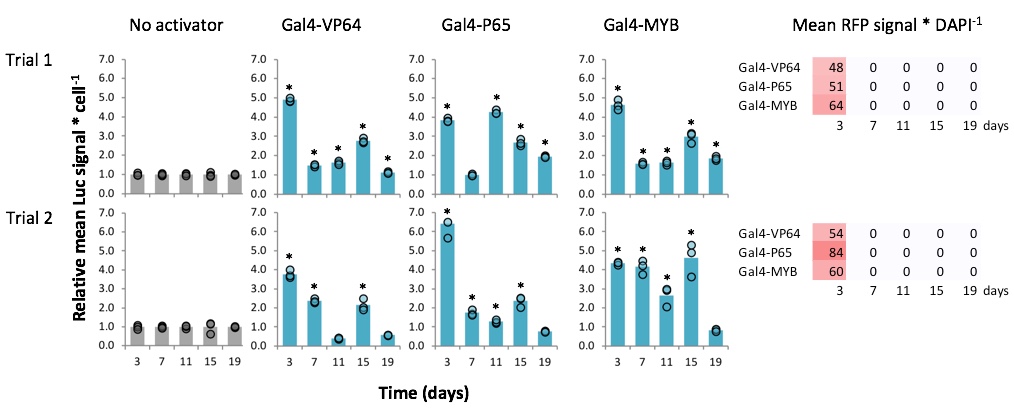

**Figure S4**. Additional trial of time course experiments with Gal4-AAP-expressing cells. Luciferase assays for Trial 2 were performed as described for Trial 1 from Figure 5A. As a high throughput alternative to flow cytometry, a microplate reader was used to measure both RFP signal and DAPI signal from Hoescht-stained cells in 96-well plates *in situ*. The assay and data analysis were performed as we have described previously [S14]. Normalized RFP signal values are shown in each table (right). Asterisks (*) = *p* < 0.05 for mean values greater than the mean for the “No activator” negative control sample.

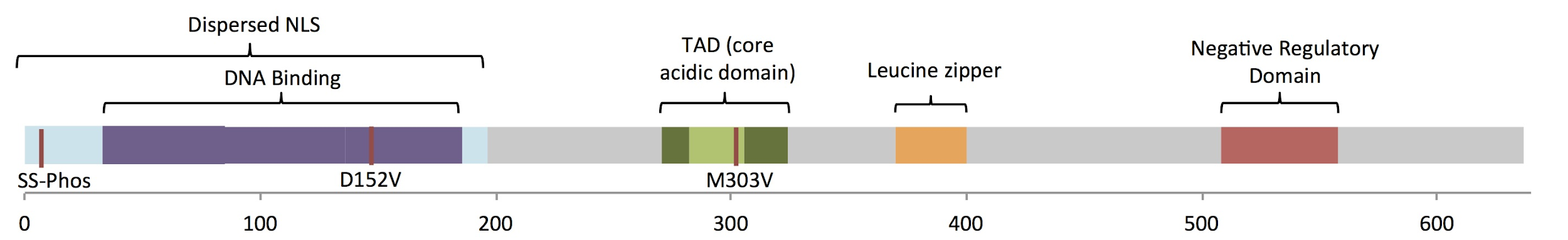

**Figure S5**. Annotated motifs within the full-length MYB protein (AAA52032.1). From left to right: Dispersed nuclear localization signal (NLS) (M1-Q200; in light blue); Casein kinase II phosphorylation sites that reduce MYB DNA binding when phosphorylated and D152V mutation site that nullifies limited MYB pioneer function by disrupting DNA binding site recognition (S11, S12, D152; in red) [S15]; Repeat regions (R1-R3) that facilitate MYB binding to its recognition element (G34-L86, N87-L138, N139-M189; in purple) [S16, S17]; Transcription activation domain (P275-W327; in green) [S16]; Core acidic domain of TAD that facilitates interaction with CBP/p300 (D286-L309; in bright green) [S18]; M303V mutation that disrupts p300 recruitment and thus, activation by MYB (M303; in red) [S18, S19]; Leucine zipper domain that interacts with other cellular proteins (L383-L403; in orange) [S18]; Negative regulatory domain (V512-P566; in red), removal of which increases MYB-mediated expression increase [S20].

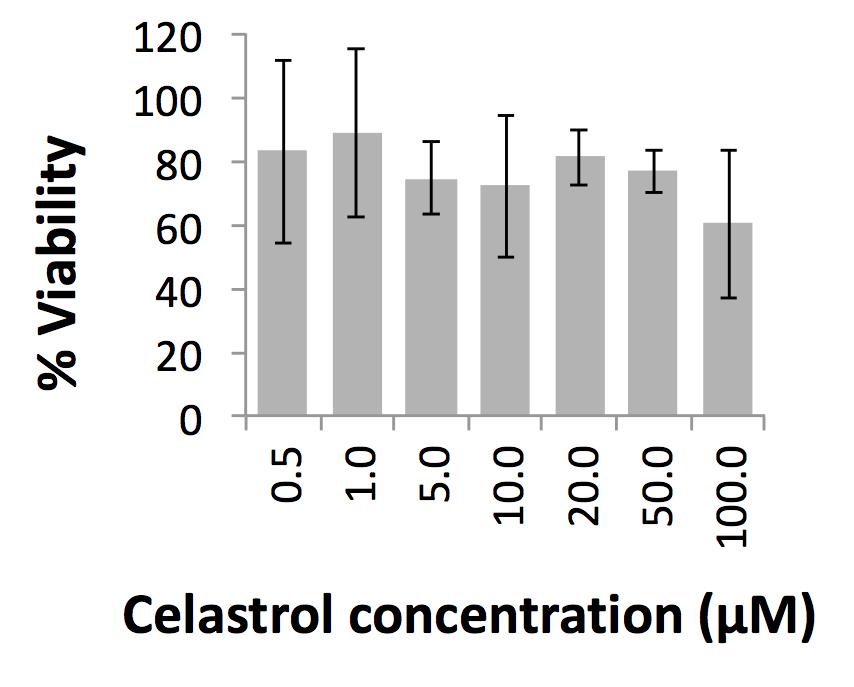

**Figure S6**. MTT cell viability assays. An MTT ((3‐(4,5‐Dimethylthiazol‐2‐yl)−2,5‐diphenyltetrazolium bromide)) cell viability assay to determine the effects of celastrol on cell survival was performed as described previously by Godeshala et al. [S21] In brief, Gal4-EED/luc cells were treated with celastrol diluted at different concentrations in Gibco DMEM high glucose. Cells were incubated with the drugs for six hours before being washed and cultured for 3 hours in drug free medium containing MTT reagent solution. Finally, cells were incubated with methanol:dimethyl sulfoxide (1:1) at room temperature for 30 minutes and mixed thoroughly before absorbance was read at 570 nm. The relative cell viability (%) was calculated from [ab]test/[ab]control × 100%. For each sample, the final absorbance/cell viability was reported as the average of values measured from three wells in parallel. Bars = standard error. We found no significant impact on cell viability after treatment with celastrol at any of the concentrations tested.

**SUPPLEMENTAL REFERENCES**

1. Beerli RR, Segal DJ, Dreier B, Barbas CF 3rd. Toward controlling gene expression at will: specific regulation of the erbB-2/HER-2 promoter by using polydactyl zinc finger proteins constructed from modular building blocks. Proc Natl Acad Sci U S A. 1998;95:14628–33.

2. Liu PQ, Rebar EJ, Zhang L, Liu Q, Jamieson AC, Liang Y, et al. Regulation of an endogenous locus using a panel of designed zinc finger proteins targeted to accessible chromatin regions. Activation of vascular endothelial growth factor A. J Biol Chem. 2001;276:11323–34.

3. Zobel A, Kalkbrenner F, Vorbrueggen G, Moelling K. Transactivation of the human c-myc gene by c-Myb. Biochem Biophys Res Commun. 1992;186:715–22

4. Kawasaki H, Schiltz L, Chiu R, Itakura K, Taira K, Nakatani Y, et al. ATF-2 has intrinsic histone acetyltransferase activity which is modulated by phosphorylation. Nature. 2000;405:195–200.

5. Yang L, Wang H, Luo X, Mao P, Tian W, Shi Y, et al. Virion protein 16 induces demethylation of DNA integrated within chromatin in a novel mammalian cell model. Acta Biochim Biophys Sin . 2011;44:154–61.

6. Milne TA, Briggs SD, Brock HW, Martin ME, Gibbs D, Allis CD, et al. MLL targets SET domain methyltransferase activity to Hox gene promoters. Mol Cell. 2002;10:1107–17.

7. Nishioka K, Chuikov S, Sarma K, Erdjument-Bromage H, Allis CD, Tempst P, et al. Set9, a novel histone H3 methyltransferase that facilitates transcription by precluding histone tail modifications required for heterochromatin formation. Genes Dev. 2002;16:479–89.

8. Antonysamy S, Bonday Z, Campbell RM, Doyle B, Druzina Z, Gheyi T, et al. Crystal structure of the human PRMT5:MEP50 complex. Proc Natl Acad Sci U S A. 2012;109:17960–5.

9. Serandour AA, Avner S, Percevault F, Demay F, Bizot M, Lucchetti-Miganeh C, et al. Epigenetic switch involved in activation of pioneer factor FOXA1-dependent enhancers. Genome Res. 2011;21:555–65.

10. Clark KL, Halay ED, Lai E, Burley SK. Co-crystal structure of the HNF-3/fork head DNA-recognition motif resembles histone H5. Nature. 1993;364:412–20.

11. Wang W, Côté J, Xue Y, Zhou S, Khavari PA, Biggar SR, et al. Purification and biochemical heterogeneity of the mammalian SWI-SNF complex. EMBO J. 1996;15:5370–82.

12. Daer RM, Cutts JP, Brafman DA, Haynes KA. The Impact of Chromatin Dynamics on Cas9-Mediated Genome Editing in Human Cells. ACS Synth Biol. 2017;6:428–38.

13. Hansen KH, Bracken AP, Pasini D, Dietrich N, Gehani SS, Monrad A, et al. A model for transmission of the H3K27me3 epigenetic mark. Nat Cell Biol. 2008;10:1291–300.

14. Tekel SJ, Barrett C, Vargas D, Haynes KA. Design, Construction, and Validation of Histone-Binding Effectors in Vitro and in Cells. Biochemistry. 2018;57:4707–16.

15. Fuglerud BM, Lemma RB, Wanichawan P, Sundaram AYM, Eskeland R, Gabrielsen OS. A c-Myb mutant causes deregulated differentiation due to impaired histone binding and abrogated pioneer factor function. Nucleic Acids Res. 2017;45:7681–96.

16. George OL, Ness SA. Situational awareness: regulation of the myb transcription factor in differentiation, the cell cycle and oncogenesis. Cancers . 2014;6:2049–71.

17. Joaquin M, Watson RJ. Cell cycle regulation by the B-Myb transcription factor. Cell Mol Life Sci. 2003;60:2389–401.

18. Wang D-M, Lipsick JS. Mutational analysis of the transcriptional activation domains of v-Myb. Oncogene. 2002;21:1611–5.

19. Sandberg ML, Sutton SE, Pletcher MT, Wiltshire T, Tarantino LM, Hogenesch JB, et al. c-Myb and p300 regulate hematopoietic stem cell proliferation and differentiation. Dev Cell. 2005;8:153–66.

20. Gonda TJ, Favier D, Ferrao P, Macmillan EM, Simpson R, Tavner F. The c-Myb Negative Regulatory Domain. In: Current Topics in Microbiology and Immunology. 1996. p. 99–109.

21. Godeshala S, Nitiyanandan R, Thompson B, Goklany S, Nielsen DR, Rege K. Folate receptor-targeted aminoglycoside-derived polymers for transgene expression in cancer cells. Bioeng Transl Med. 2016;1:220–31.
